# During an inflammatory response, zebrafish *tnfa* and *tnfb* are expressed by different cell types and have distinct expression kinetics

**DOI:** 10.1101/2025.11.04.686030

**Authors:** Kaylee S.E. van Dijk, Christina Begon-Pescia, Boudewijn A. de Bruin, Resul Özbilgiç, Philip M. Elks, Mai E. Nguyen-Chi, Maria Forlenza

## Abstract

Tumour necrosis factor (TNF, TNFSF2) is a central regulator of inflammation. While mammalian TNF biology is well understood, its pleiotropic role in other vertebrates remains unclear. Zebrafish (*Danio rerio*) possess two TNF homologues, *tnfa* and *tnfb*. Previous studies have focused mainly on *tnfa*, leaving *tnfb* largely unexplored. We generated two transgenic reporter lines, *Tg(−3.2tnfb:eGFP-F)^ump21Tg^* and *Tg(−6.2tnfb:mCherry-F)^ump22Tg^*, to track *tnfb* expression. Combining these reporter lines with high-resolution *in vivo* microscopy, other reporter lines, and public scRNAseq datasets, we define, in a cell type-specific manner, *tnfb* expression kinetics compared to those of *tnfa* and *il1b*. We found constitutive *tnfb* expression in neuromast mantle cells from 32 hpf, and inducible expression in macrophage and neutrophil subsets following caudal fin fold amputation or *E. coli* injection. After wounding, *tnfb* expression appeared first in macrophages, followed by neutrophils, with sequential *tnfb*, *tnfa*, and *il1b* expression in macrophages. These findings reveal distinct *tnfb* dynamics compared to *tnfa*, suggesting differential roles of these two cytokines, and establishing zebrafish as a model to study the pleiotropic functions of Tnf in vertebrates.

## Introduction

In mammals, tumour necrosis factor (TNF, previously TNF-α (alpha); TNFSF2 genenames.org) is a key orchestrator of inflammation, among other processes, and is expressed by and has effect on several cell types, including immune cells. The main producers of and responders to TNF are monocytes and macrophages (Aggarwal, 2003; Beutler and Ceramit, 1989; Dubravec et al., 1990). When a cell responds to TNF signalling, an intracellular signalling cascade is activated leading to a broad range of cellular activities, including proliferation, survival, differentiation, immune mediators production or cell death (Baud and Karin, 2001; Horiuchi et al., 2010; Preedy et al., 2024). These seemingly opposite activities are essential for maintaining a delicate balance between protective inflammatory responses and excessive tissue damage. In fact, numerous studies describe how a tight regulation of TNF during the early phase of inflammation is required to control inflammation or infection, as opposed to pathology related to uncontrolled TNF expression and subsequent exacerbated innate immune responses (Aggarwal, 2003; Aggarwal et al., 2012; Beutler and Ceramit, 1989; Dubravec et al., 1990; Feldmann and Maini, 2003).

Despite the substantial progress been made in the characterization of mammalian TNF biology, the pleotropic nature of TNF during inflammation or infection in other vertebrate species remains elusive. Interestingly, while in the genome of mammals, birds, reptiles and amphibians, only one copy of *TNF* has been identified (Secombes et al., 2016), in the majority of bony fish (teleost) genomes analysed so far, two or more *tnf* homologues can be identified (Cui et al., 2020; Grayfer et al., 2008; Hong et al., 2013; Kadowaki et al., 2009; Kajungiro et al., 2015; Y. Li et al., 2021; Milne et al., 2017; Nascimento et al., 2007; Pleić et al., 2015; Saeij et al., 2003). They have been subdivided into two groups based on distinct structural and functional properties (Hong et al., 2013). Zebrafish (*Danio rerio*) possess two TNF homologs (Kinoshita et al., 2014), *tnfa* belonging to group II located on chromosome 19, and *tnfb* belonging to group I located on chromosome 15 (Hong et al., 2013), and will be referred to as *tnfa* and *tnfb,* or *tnf* when referring to both. According to zebrafish nomenclature conventions, homologous genes are assigned suffixes a or b, and these should not be confounded with the designations alpha or beta. Thus, tnfa is not synonymous of TNFα and tnfb is not synonym of TNFβ (also referred to as lymphotoxin-alpha (LTA)). While *tnfa* has been extensively studied in the context of wounding or regeneration (Keatinge et al., 2021; Miskolci et al., 2019; Nelson et al., 2013; Nguyen-Chi et al., 2017, 2015; Tsarouchas et al., 2018), and during various bacterial infections (Bernut et al., 2016; Kam et al., 2022; Lewis and Elks, 2019; Miskolci et al., 2019), *tnfb* may have important roles in development and inflammation, yet has remained largely unexplored. Although direct functional evidence is sparse, it was suggested that *tnfb* alongside *tnfa* is involved in liver development (Qi et al., 2010). In the context of infection-induced inflammation, zebrafish larvae pre-injected with anti-Tnfb antibodies showed increased survival after *Vibrio vulnificus* infection (Li et al., 2021), suggesting that regulation of Tnfb expression may determine susceptibility or resistance to the infection. Whether zebrafish Tnfa (group II) and Tnfb (group I) have overlapping as well as distinct roles during inflammation or infection, and how these relate to the better-known pleiotropic functions of mammalian TNF, is a knowledge gap that awaits further investigation. Zebrafish are well-studied model organisms and offer several advantages, including larval transparency, a well annotated genome amenable to genetic manipulation, availability of several transgenic lines marking innate immune cells and cytokine-expressing cells (Benard et al., 2015; Bertrand et al., 2010; Dóró et al., 2019; Ellett et al., 2011; Jacobs et al., 2021; Lawson and Weinstein, 2002; Page et al., 2013; Petrie-Hanson et al., 2009; Renshaw et al., 2006; White et al., 2008), altogether making them suited to study the role of *tnfb* during inflammation. Furthermore, larval zebrafish lack a mature adaptive immune system for the first 2-3 weeks of development (Lam et al., 2004; Langenau et al., 2004; Torraca and Mostowy, 2018), offering the opportunity to focus on inflammatory responses driven by innate immune cells, especially macrophages and neutrophils.

For Tnfa and Il1b, a proinflammatory role and association with inflammatory macrophages have been established in the context of wound healing and bacterial challenges (Hasegawa et al., 2017; Lewis and Elks, 2019; Nguyen-Chi et al., 2017, 2014; Ogryzko et al., 2019). *tnfa* positive macrophages were shown to be essential during wound healing (Nguyen-Chi et al., 2017), to be highly phagocytic and to exert antimicrobial activities (Bernut et al., 2016; Clay et al., 2008; Nguyen-Chi et al., 2015). In such inflammatory contexts, also neutrophils express *il1b* alongside macrophages (Ogryzko et al., 2019). The role of Tnfb however, remains unclear. Previous studies that investigated *tnfb* gene expression, showed that it was enriched in macrophages during the proinflammatory phase of the response to caudal fin fold amputation (Nguyen-Chi et al., 2015). However, which cell-types express *tnfb,* its constitutive expression, and expression kinetics during inflammation *in vivo,* are largely unexplored.

In this study, we generated and characterized novel promotor-driven transgenic lines marking *tnfb-*expressing cells. We focused on the early phase of inflammation triggered by two well-established inflammation models: wounding, after caudal fin fold amputation (Begon-Pescia et al., 2022; Nguyen-Chi et al., 2017, 2015; Sipka et al., 2022, 2021), or infection, after *Escherichia coli* (*E. coli*) injection (Nguyen-Chi et al., 2015), to define, in a cell type-specific manner, the expression kinetics of *tnfb* compared to those of *tnfa* and *il1b*. We report that mantle cells of neuromast constitutively express *tnfb* from 32 hours post-fertilization onwards. After injury, we show that a subset of macrophages expressed both, *tnfa* and *tnfb*, whereas a subpopulation of neutrophils expressed *tnfb* and not *tnfa*. Interestingly, after wounding, *tnfb* expression kinetics differ in macrophages and neutrophils, with *tnfb* expression detected in macrophages preceding its detection in neutrophils. When analysing the kinetics of expression of *tnfb, tnfa* and *il1b* after wounding, our data indicate that, at least in macrophages, a sequential expression occurs, with *tnfb* expressed first followed by *tnfa* and *il1b*. Altogether, we demonstrate that *tnfb* is expressed by various cell types and has distinct expression patterns from *tnfa*, and propose that by studying both TNF homologues, the zebrafish model is particularly suited to dissect the pleiotropic nature of TNF during inflammation and infections.

## Material and methods

### Zebrafish line and maintenance

Zebrafish were kept and handled according to standard protocols (Westerfield, 2000; zfin.org) and local animal welfare regulations of The Netherlands. Zebrafish larvae were raised in E3 medium (4.96 mM NaCl, 0.18 mM KCl, 0.44 mM CaCl_2_ and 0.40 mM MgCl_2_·6H_2_O in demi water at pH 7.2) at 28°C.

The following zebrafish lines were used in this study: transgenic *Tg(mpx:eGFP)^i114^* (Renshaw et al., 2006), *Tg(mpeg1:eGFP)^gl22^* (Ellett et al., 2011)*, Tg(mpeg1:mCherry-F)^ump2Tg^* (Nguyen-Chi et al., 2014)*, TgBAC(il-1b:eGFP)*^sh445^ (Ogryzko et al., 2019) referred to as Tg(il1b:eGFP)*, Tg(tnfa:eGFP-F)^ump5Tg^* (Nguyen-Chi et al., 2015), or crosses thereof. Two transgenic zebrafish lines express a farnesylated (membrane-targeting sequence) mCherry (mCherry-F) or eGFP (eGFP-F) under the control of the *mpeg1.1* or *tnfa* promoter, respectively.

### *tnfb* transgenic line construction

The *tnfb* promotor (gene: ENSDARG00000013598) was amplified from zebrafish genomic DNA using primers Ztnfb-P3 FW: 5’-TAGCCAAGCCTACTTGTTGC-3’ or Ztnfb-P7 FW: 5’-AGGGTGACGATAATGTCAAC-3’ and Ztnfb_E1N RV: 5’-TAATAGCGGCCGCTTTCGTATCTCACCATGCTG-3’. The resulting fragments were phosphorylated using T4PNK, digested with *NotI* and cloned in a derivative of pBSKI2 vector, upstream of either farnesylated eGFP (eGFP-F) (Thermes et al., 2002) or mCherry (mCherry-F). The resulting plasmids harbour either a 3.2-kb or 6.2-kb fragment of the zebrafish *tnfb* promoter, and each include part of the first coding exon using the endogenous ATG codon of *tnfb* to drive the translation of eGFP-F or mCherry-F. The expression cassette is flanked by two *I-SceI* sites. Either of these constructs was co-injected in fertilized eggs with the enzyme *I-SceI* (New England Biolabs, R0694S). Developing embryos underwent caudal fin fold amputation at 3 dpf (days post-fertilization), and those with clearly visible green or red fluorescence were raised as putative founders. Their offspring was tested in a similar manner to establish the stable transgenic lines *Tg(−3.2tnfb:eGFP-F)^ump21Tg^* and *Tg(−6.2tnfb:mCherry-F)^ump22Tg^,* referred to as *Tg(tnfb:eGFP-F)* and *Tgtnfb:mCherry-F)*.

### Caudal fin fold amputation *and E. coli* injection of zebrafish larvae

Caudal fin fold amputation was performed on 3 dpf larvae as described previously (Begon-Pescia et al., 2022). In brief, the caudal fin of anaesthetised larvae was transected with a sterile scalpel, posterior to the muscle and notochord.

For infection studies, 3 dpf larvae were injected with 2×10^3^ CFU *Escherichia coli* K12 harbouring an E2-Crimson expression plasmid, from here on referred to as *E. coli* crimson. Bacteria were grown overnight at 37 °C on Luria Broth (LB) agar plates (1.5% BactoTM Agar (BD, 214010), 1% BactoTM Trypton (GibcoTM, 211705), 0.5% BactoTM Yeast extract (GibcoTM, 212750) and 0.5% NaCl (Merck KGaA, 1.37017) in demi water) supplemented with 50 μg/ml ampicillin (A9518, Sigma-Aldrich), and 0.25 mM Isopropyl β-D-1-thiogalactopyranoside (IPTG, Promega, V395A). Bacteria were harvested from the plate, washed twice and resuspended in sterile PBS (Gibco, 10010-015) to measure the OD600 (Camspec M107 Spectrophotometer), next bacteria were resuspended in sterile 2% (w/v) polyvinylpyrrolidone in PBS (PVP, Sigma-Aldrich, PVP40-50G) solution. Anesthetized larvae were injected intramuscularly with 2×10^3^ CFU (in 1 nl) of *E. coli* crimson solution or 2% PVP alone.

### *In vivo* imaging of zebrafish

Prior to imaging, zebrafish larvae were anaesthetised in 0.017% MS-222 (ethyl 3-aminobenzoate methanesulfonate, Sigma-Aldrich, A5040-25G) and embedded in 1% UltraPure^TM^ Low Melting Point Agarose (Thermo Fisher, 16520-050) at the bottom of a 35 mm FluoroDish^TM^ (World precision instruments, 13012026). Andor-Revolution spinning disk confocal (Yokogawa) on a Nikon Ti Eclipse PFS3 microscope, 40x (1.15 NA, 0.61-0.59 mm WD) WI objective, 20x (0.75 NA, 1.0 mm WD) objective were used with the following settings: dual pass 523/561: GFP excitation: 488 nm, emission: 510-540 nm, EM gain: 150-300 ms, digitizer: 10 MHz (14-bit); RFP excitation: 561 nm; emission: 589-628 nm, EM gain: 200-450 ms, digitizer: 10 MHz (14-bit); CY5 excitation: 640 nm; emission: 665-705 nm, EM gain: 200-450 ms, digitizer: 10 MHz (14-bit); BF DIC EM gain: 180-250 ms, digitizer: 10 MHz (14-bit). Typically for time-lapse videos, Z-stacks spanning 20-60 µm at 1 µm steps were acquired every 3-18 min. For individual z-stack acquisitions the 20x or 40x objective was used, as indicated by the figure legends.

For detailed imaging of neuromasts, a Leica SP8-DIVE multiphoton confocal microscope (Leica Microsystems) equipped with a near infrared femtosecond sapphire pulse laser and a 20x (0.75 NA, 1.0 mm WD) objective was used with the following settings: GFP excitation 900 nm, mCherry excitation 1050 nm using preset GFP or mCherry detection modules available in the Leica Application Suite software for DIVE spectral multiphoton detectors. The image scan speed was 4 µs/pixel with an image size of 512 × 512 pixels.

Images were manually analysed with ImageJ-FIJI (version 2.1.0, Schindelin et al., 2012). After amputation, the number of recruited macrophages, neutrophils and cytokine-expressing cells was quantified after brightness and contrast adjustment in ImageJ-FIJI for better visualizations. Cells were counted at the site of injury, defined as a 105 μm width region centred around the posterior tip of the notochord.

For total cell fluorescence acquisition after *E. coli* injection, transgenic zebrafish were positioned on preheated flat agarose plates (1% agarose in E3 medium containing 0.017% MS-222) and imaged with a Leica M205FA fluorescence stereo microscope (Leica Microsystems). The image acquisition settings were as follows: Zoom: 58-61x, Gain: 2.5, Exposure time (ms): 300 (BF)/1000 (GFP)/6000 (mCherry)/ 6500 (Crimson), Intensity: 200 (BF)/100 (GFP)/230 (mCherry)/ 250 (Crimson), Contrast: 255/255 (BF)/ 70/255 (GFP)/ 15/255 (mCherry)/ 20/255 (Crimson). Quantification of total cell fluorescence was performed using the free-form selection tool and by accurately selecting the area around the fluorescent reporter, selected areas were saved using the ROI manager tool. At later timepoints or for PVP-injected larvae the equivalent area positioned around the site of injection. Area integrated intensity and mean grey values of each selected area were acquired. Analysis was performed using the following formula (Fitzpatrick, 2014): Corrected Total Cell Fluorescence (CTCF) = integrated density – (area x mean grey background value). The mean grey background value was the average of three random areas outside of the fish selected in each image.

### Fluorescence-activated cell sorting, isolation of mRNA and gene expression

Fluorescence-activated cell sorting (FACS) and mRNA isolation were performed as described previously (Begon-Pescia et al., 2022). Briefly, the protocol is divided in five steps: (1) caudal fin fold amputation; (2) Enzyme-free mechanical cell dissociation was performed by passing larvae through a cell strainer with a 70 µm mesh (Falcon™, 352350) using sorting buffer (0.9x Dulbecco’s Phosphate-Buffered Saline without calcium and magnesium (Life Technologies, 14190094), with 2% heat inactivated foetal bovine serum (Gibco, 10500064) and 2 mM EDTA (Invitrogen, 15575-020) kept at 4°C. To ensure a single cell suspension, cells were passed through a 40 µm mesh (Falcon™ 352340) and kept on ice until sorting; (3) sorting of eGFP^+^ and eGFP^-^cells form *Tg(tnfb:eGFP-F)* or sorting of mCherry^+^ and mCherry^-^cells from *Tg(tnfb:mCherry-F)* larvae (3 dpf, 2-3 hpa) with FACS Aria™ IIu (BD biosciences) (**Supplementary Fig 1**); (4) total RNA extraction from sorted cells using the RNeasy micro Kit (Qiagen, 74004); (5) cDNA synthesis from 10 ng of total RNA using M-MLV Reverse Transcriptase Transcription kit (Invitrogen, 28025-013). Real-time quantitative PCR (RT-qPCR) was performed on a LightCycler 480 system (Roche) using SYBR Green (*SensiFAST™ SYBR*^®^ No-ROX Kit, 98050) as reaction mix according to the manufacturers instruction and primers listed in **table 1**. Program: 5 min at 95°C, 45 cycles of denaturation 15 s at 95°C, annealing 10 s at 64°C, and elongation 20 s at 72°C. Expression levels were determined with the LightCycler analysis software (version-1.5.1). The relative expression level of a given mRNA was calculated using the formula 2^−ΔCt^ with housekeeping gene *eef1a1l1* (previously *ef1a*) as internal reference.

**Table 1.**
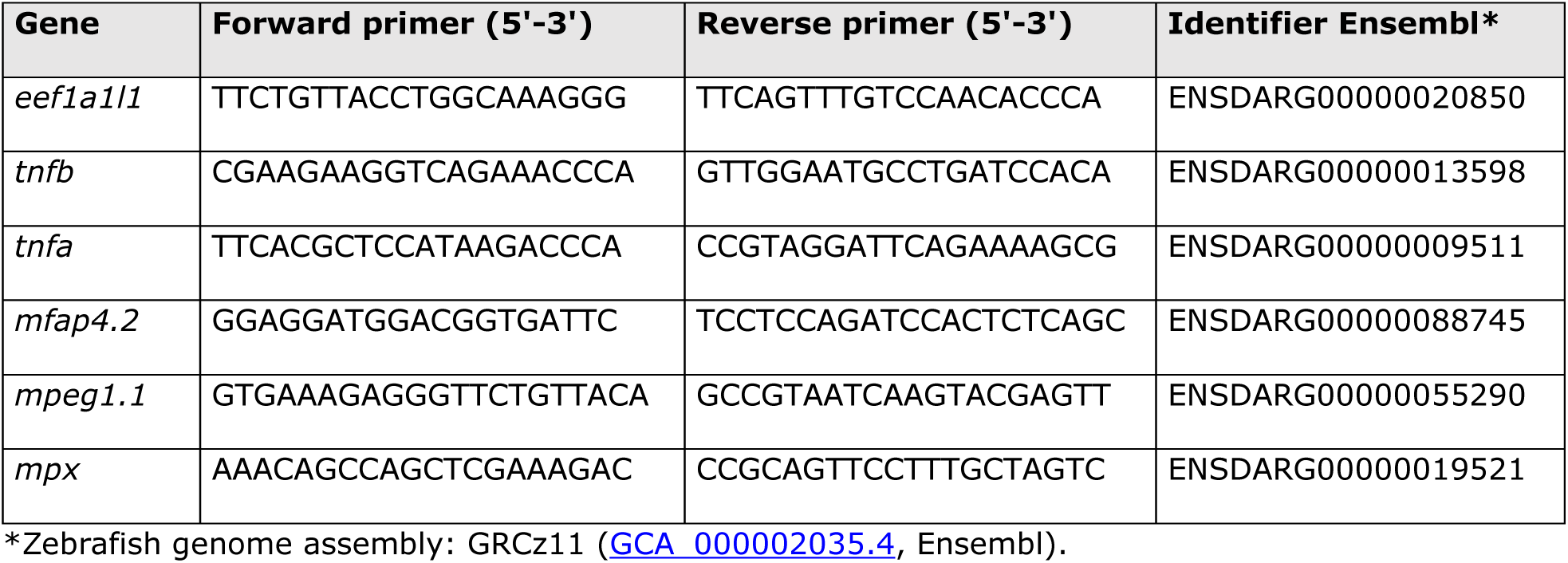
List of RT-qPCR primers used in this study.

### Whole-mount *in situ* hybridization

A *tnfb* probe was amplified from total cDNA derived from 3 dpf zebrafish larvae. The following primers were used for PCR: Forward 1_*tnfb* 5’-GCAGCATGGTGAGATACGAA-3’ and Reverse 1_*tnfb* 5’-ACTCTCACTGCATCGGCTTT-3’. The 890 bp amplicon was ligated into a pCRII-TOPO plasmid (TOPO™ TA Cloning™ Kit, Invitrogen, 45-0640). Agarose-gel electrophoresis analysis was performed to confirm the size of the insert by digesting the *tnfb-*pCRII-TOPO plasmid with *EcoRI*-HF (New England Biolabs, R3101S), which recognition sites flank the PCR insert. Sequencing was performed by Eurofins genetics using their standard SP6 primer: 5’-CATTTAGGTGACACTATAG-3’. Similarly, a 786 nt *mpx* probe was generated using primers: Forward 1_*mpx* 5’-AAACAGCCAGCTCGAAAGAC-3’ and Reverse 1_*mpx*: 5’-CCGCAGTTCCTTTGCTAG-3’.

To generate RNA probes, the plasmids were linearized by restriction digestion with *HindIII* (Thermo Scentific, FD0504) or *NotI-FD* (Thermo Scentific, FD0593) to facilitate *in vitro* transcription of antisense or sense digoxigenin (DIG)-labelled (Roche, 11277073910) probes, using T7 or Sp6 RNA polymerase (Promega, T7: M0251S or Sp6: P1085 RNA polymerase kit) according to the manufacturer’s instruction. T7 promotor was used to generate antisense probe for *tnfb*, whereas SP6 promotor was used to generate sense probe for *tnfb* and antisense probe for *mpx*. Probe integrity was confirmed by agarose-gel electrophoresis.

Larvae were euthanized with an overdose of anaesthetic (0.4% MS-222), and fixed overnight in 4% paraformaldehyde (PFA) at 4°C, followed by three washes with phosphate-buffered saline (PBS) with 0.1% (v/v) tween 20 (PBT), followed by gradual dehydration by 5 min incubations in increasing concentrations of methanol (25%, 50%, 75%, 100%). Larvae were stored in 100% methanol at −20°C for at least one week. Larvae were gradually re-hydrated using decreasing concentrations of methanol (75%, 50%, 25%), followed by three washes in PBT. In *situ hybridization* on whole-mount larvae was performed as previously described (Thisse and Thisse, 2008), with adjusted enzymatic digestion of larvae. To preserve tissue structure of neuromasts a mild enzymatic digestion, with 2 μg/mL *Proteinase K* (Promega, MC5005) for 17 min at room temperature (21°C), was performed (Steiner et al., 2014). Digestion was stopped by 20 min incubation in 4% paraformaldehyde (PFA) at room temperature, followed by three washes in PBT, and 6h incubation in hybridization buffer (nuclease free water containing 50% formamide (Sigma, S417117), 5x saline-sodium citrate (SSC, Sigma, 118K8403), 500 µg/ml tRNA (Sigma, R8759_500UN), 50 µg/ml heparin (Sigma, H3393-50KU), 0.1% tween, and 1M citric acid (Sigma, 71402) to adjusted to pH 6). Prior to hybridization of larvae, the probe was linearized by 3 min incubation at 80°C, directly followed by 5 min incubation on ice. Overnight hybridization was at 65°C, per ten larvae 150 ng linearized probe in 200 µl was used.

Several stringent washing steps were performed at 65°C using freshly prepared solutions, containing various concentration of formamide, SSC and tween (as described step-by-step by Thisse and Thisse, 2008). Hereafter, larvae were incubated in blocking buffer (PBT with 2% normal sheep serum (Roche, 013-000-121) and 2 mg/ml Bovine Serum Albumin (Sigma, A3059) for 5h at room temperature. The transcript’s expression pattern was visualized using anti-DIG antibody conjugated to alkaline phosphatase (1:500, Roche, 11093274910) overnight at 4°C in the dark, followed by six washed in PBT for 15 min each. Next a series of four stringent washed in freshly prepared NTMT (Final concentration: 100 mM NaCl, 100 mM Tris-HCL pH 9.5, 50 mM MgCl2, 0.1% tween in nuclease free water), followed by incubation in nitro blue tetrazolium/5-bromo-4-chloro-3-indolyl phosphate substrate (NBT/BCIP, Roche, 11681451001) until sufficient colour developed. Reactions were terminated by washing larvae in PBT and incubation in 4% PFA for 30 minutes. Incubation with *tnfb* sense probe was carried out in parallel as a negative control, and *mpx* antisense probe hybridization was carried out as a positive control. Prior to imaging, larvae were cleared sequentially: larvae were incubated for five minutes in increasing concentrations of ethanol (30%, 50%, 70%) in PBS and incubated twice in 100% ethanol. Finaly, larvae were incubated in increasing concentrations of glycerol (30%, 50%, 80%) in PBS.

Images were acquired using a Zeiss Stemi 508 stereo microscope (Zeiss Microscopy), equipped with Moticam 10+ 10.0 MP colour camera (Motic) and Motic Images Plus software (version 3.0 ML). Alternatively, images were acquired using Leica DM6b automated upright digital microscope (Leica Microsystems), controlled by Leica LASX software (version 3.6.0.), equipped with a 20x (NA 0.8, DIC) short distance objective and DFC450C CCD colour camera (Leica Microsystems).

### Exploration of public single-cell RNA sequencing datasets

This paper used publicly available single-cell RNA sequencing (scRNAseq) datasets, explored through their respective web-based visualization platforms: I) data of homeostatic neuromasts retrieved from <u>Zebrafish Neuromast scRNAseq</u> (Lush et al., 2019), II) data from neuromasts at multiple timepoints after hair cell ablation retrieved from <u>Neuromastregeneration scRNAseq</u> (Baek et al. 2022), and III) data of multiple zebrafish developmental stages (3 – 120 hpf) retrieved from <u>Daniocell</u> database (Farrell et al., 2018; Sur et al., 2023). the latter, was explored using the companion application <u>Daniocell Desktop</u> (v1.0.3, Evans and Farrell, 2025). Each dataset provided pre-processed and annotated scRNAseq data, allowing exploration of gene expression across defined cell clusters.

### Statistical analysis

Statistical analyses of gene expression, cell number and total fluorescence data were performed in GraphPad PRISM 10 (GraphPad Software). The specific statistical tests used are indicated per figure legend. In all cases, \**p*<0.05 was considered significant.

## Results

### *tnfb* is constitutively expressed in neuromast cells during early development

In the present study, two transgenic reporter lines marking *tnfb* were generated: one in which a 3.2-kb portion of the predicted *tnfb* promotor drives farnesylated eGFP (eGFP-F) expression and a second in which a 6.2-kb portion of the predicted *tnfb* promotor drives farnesylated mCherry (mCherry-F) expression. When the constructs were independently injected into wildtype embryos, reporter expression was observed along the lateral line (in neuromast cells) of F0, suggesting that both promotors could drive fluorescent protein expression in unchallenged conditions. Five stable transgenic zebrafish lines (three *Tg(tnfb:eGFP-F)* and two *Tg(tnfb:mCherry-F)*) were recovered that retained the fluorescent pattern observed in neuromasts. Due to higher fecundity in *Tg(−3.2tnfb:eGFP-F)^ump21Tg^* and *Tg(−6.2tnfb:mCherry-F)^ump22Tg^*, these two lines were used hereafter. In crosses of *Tg(tnfb:eGFP-F);Tg(tnfb:mCherry-F)* transgenic lines, earliest expression of both transgenes was detected concomitantly around 32 hpf (**Fig 1A**). From 3 dpf onward, eGFP-F or mCherry-F signal was visible in the neuromasts along the entire length of the lateral line (**Fig 1B-C**). Further characterization of both transgenic lines revealed that fluorescent protein expression recapitulated the pattern of basal *tnfb* expression in 3 dpf larvae as assessed by *in situ* hybridization (**Fig 1D-E (upper panels), Supplementary Fig 2**), confirming constitutive expression of *tnfb* mRNA in neuromasts during development (**Fig 1E**). *In situ* hybridization using an antisense probe targeting *myeloperoxidase* (*mpx*) mRNA was used as a positive control for immune cell staining as it marks a population of neutrophils present in the caudal haematopoietic tissue (CHT) (**Fig 1D-E (lower panel), Supplementary Fig 2**).

**Figure 1.**
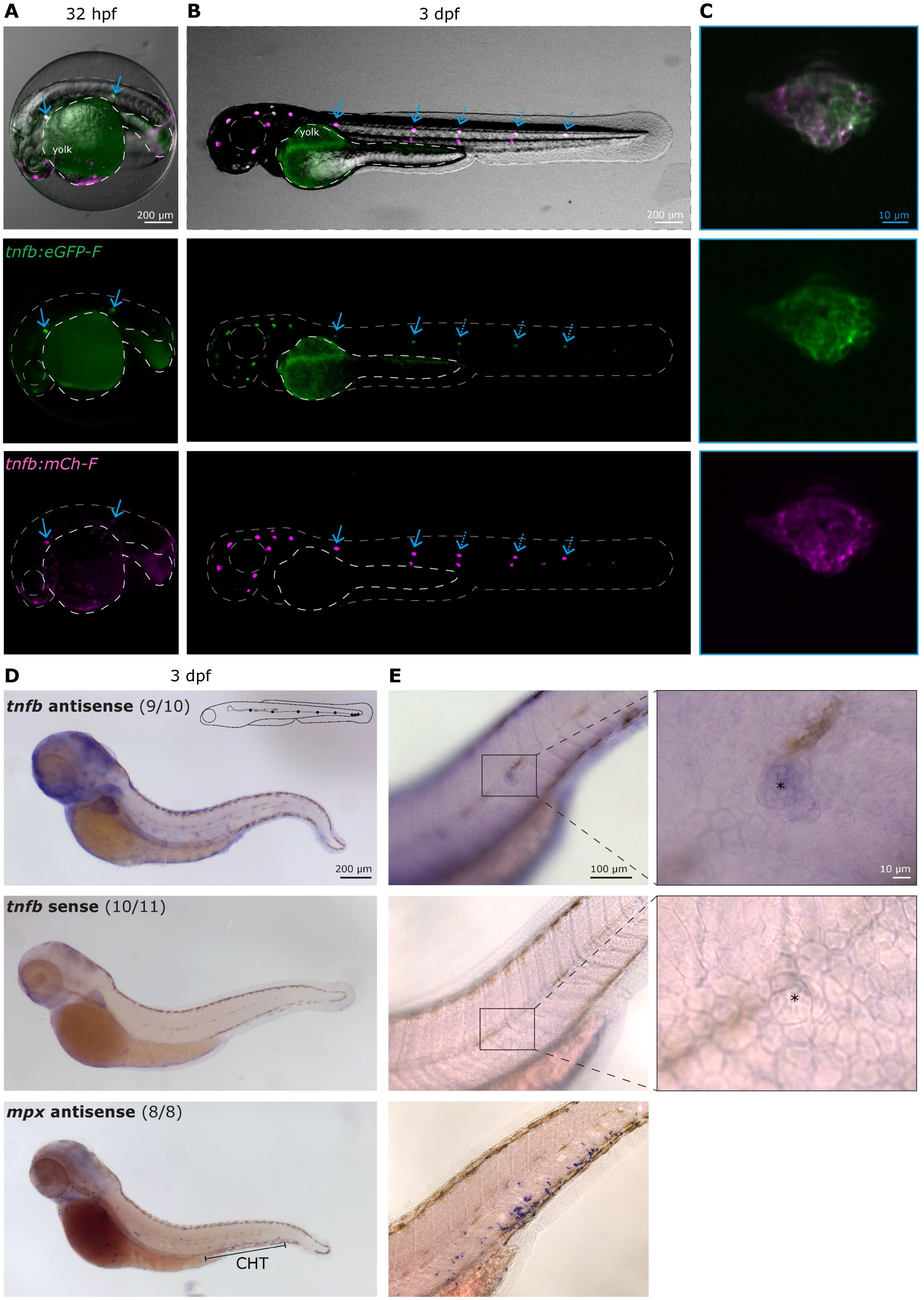
*tnfb* is constitutively expressed in neuromasts during early development. *Tg(tnfb:eGFP-F);Tg(tnfb:mCherry-F)* embryos were imaged during development using a Leica M205 stereo microscope. **A)** Earliest expression of transgenes was observed in neuromast (cyan arrows) at 32 hpf. The lateral line at this stage reaches to about halfway the length of the larvae. In some individuals, transgene-expressing cells were also observed in the extra-embryonic yolk sac. **B)** From 3 dpf onwards, transgene-expressing cells are visible in the neuromast along the lateral line, which at this stage extends to the tip of the tail. **C)** Representative maximum projection of a neuromast imaged with an Andor spinning disk confocal microscope at a 20x magnification. **D)** Representative images of zebrafish larvae acquired with a Zeiss Stemi 508 stereo microscope after wholemount *in situ* hybridisation of *tnfb* antisense (top panel), *tnfb* sense (negative control, middle panel) and *mpx* antisense probes (positive control, lower panel). Numbers following the probe name, indicate the number of larvae showing the depictured expression pattern and the total number of larvae analysed per condition. *tnfb* mRNA expression was detected in posterior neuromasts, matching the schematic drawing showing lateral neuromasts position in 3dpf zebrafish. *tnfb* sense probe showed no specific signal. *mpx* mRNA expression was mainly observed in the caudal hematopoietic tissue (CHT). **E)** Representative images of the trunk region from larvae processed in the same experiment as in D, reimaged using a Leica DMb6 microscope using a 20x magnification. Top panel: *tnfb* mRNA expression in neuromast, also visible in the inset focusing on a single neuromast (right). Middle panel: sense probe (negative control) showing no signal, confirming specificity of *tnfb* mRNA expression in neuromast. (*) indicates the position of hair cells in the centre of the neuromast. Lower panel: *mpx* mRNA expression in the CHT.

Neuromasts are organs of the lateral line, each consisting of approximately 60 cells and of a combination of different cell types (Lush et al., 2019). We investigated which sub-population(s) of neuromast cells expressed *tnfb:eGFP-F* or *tnfb:mCherry-F* by multiphoton microscopy. Expression was observed on the outer perimeter of the neuromast in both transgenics, in a dome-like or volcano-like shape typical of mantle cells (**Fig 2, Supplementary Video 1**). The cells in the core and top of the dome-like structure, composed of support and sensory hair cells, which extend through to the top of the dome, did not show *tnfb:eGFP-F* nor *tnfb:mCh-F* expression (**Fig 2, Supplementary Video 1**). The specific expression of *tnfb:eGFP-F* and *tnfb:mCh-F* in mantle cells agrees with data retrieved from two publicly available datasets reporting on RNA expression in single cells of homeostatic neuromast. There, *tnfb* is a distinguishing marker of mantle cells ((Lush et al., 2019;Baek et al., 2022), **Supplementary Fig 3**). In addition to developmental expression, a transient increase in *tnfb* mRNA expression is observed following hair cell ablation ((Baek et al. 2022), **Supplementary Fig 3D-F**). This suggest that expression is not only developmentally regulated, but also responsive to wounding. Altogether these data suggest that both newly generated transgenic lines show a similar pattern of *tnfb:eGFP-F* and *tnfb:mCh-F* expression during development, that *tnfb* is constitutively expressed in neuromast mantle cells, and that it can be further induced in response to wounding.

**Figure 2.**
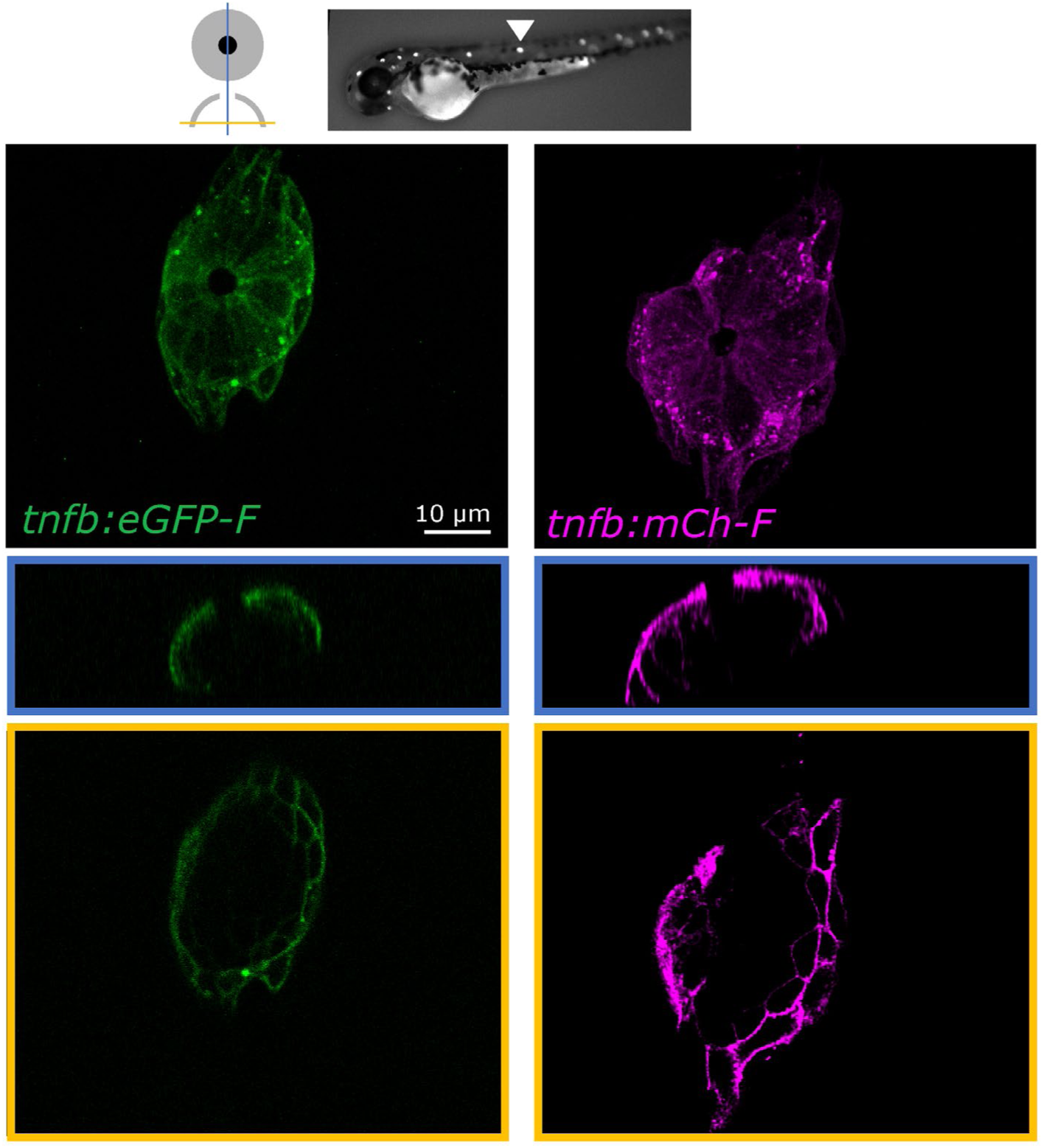
Mantle cells on the outer perimeter of neuromasts express *tnfb:eGFP-F* and *tnfb:mCherry-F*. Neuromasts of either a *Tg(tnfb:eGFP-F)* (left panel) or *of a Tg(tnfb:mCherry-F)* (right panel) transgenic zebrafish larva (3 dpf), were imaged with a Leica SP8-DIVE multiphoton microscope, at 20x magnification and 5x digital zoom. Top row: maximum projection of a neuromast in the trunk region (white arrowhead), showing *tnfb:eGFP-F* or *tnfb:mCh-F* expression localized on the outer perimeter of the neuromast, where transgene-expressing cells form a characteristic dome- or volcano-like shape, typical of mantle cells. Middle row (blue): orthogonal view, further detailing that transgene-expressing cells form a dome-like structure along the outer perimeter of neuromasts. Cells in the core and the top of the dome are *tnfb:eGFP-F* or *tnfb:mCh-F* negative. Bottom row (yellow): Single slice through the base of the neuromast, emphasizing a ring of *tnfb:eGFP-F* or *tnfb:mCh-F* positive cells around the neuromast, and the lack of transgene expression in the sensory and support cells at the centre of the dome-like structure.

### *tnfb* is transiently induced after caudal fin fold amputation

After having assessed that both transgenic lines mark constitutive *tnfb* expression in mantle cells of neuromasts, next, we assessed whether *tnfb* expression could be triggered in other cells during inflammation induced after caudal fin fold amputation. Using this wounding method, in both transgenic lines (*Tg(tnfb:eGFP-F)* and *Tg(tnfb:mCherry-F)*) or crosses thereof, cells expressing the transgene migrated towards or started to express the transgene near the site of injury (**Fig 3A, Supplementary Video 2**), matching the *in situ* hybridization pattern observed, showing *tnfb-*expressing cells localized at the wound site (**Fig 3B, Supplementary Fig 4**). Upon closer inspection, we found that cells at the wound site (within the white box) were positive for both reporter proteins (eGFP^+^mCherry^+^), while some cells were eGFP^+^mCherry^-^, eGFP^-^mCherry^+^ were never observed. Based on time-laps acquisitions we recorded cell motility over a period of 10 hours post-amputation (hpa); eGFP^+^mCherry^+^ cells were highly motile and migrated towards the wound, while in comparison, eGFP^+^mCherry^-^cells showed minimal motility, remaining stationary while undergoing occasional changes in cell shape, which remained epithelial-like (**Supplementary Video 2**). Next, the number of cells expressing the transgene(s) at the wound site was quantified at different timepoints after amputation of *Tg(tnfb:eGFP-F;tnfb:mCherry-F)* larvae (**Fig 3C-D**). Timelapse microscopy over a period of 10h (**Fig 3C**), showed that in intact fins, the number of eGFP^+^mCherry^+^ or eGFP^+^mCherry^-^cells remained constant at all investigated time points, while eGFP^-^mCherry^+^ cells were never observed. In amputated fins, the number of eGFP^+^mCherry^+^ cells increased significantly from 2h onwards compared to the same cell type in the intact group, while the number of eGFP^+^mCherry^-^cells did not change. Again, no eGFP^-^mCherry^+^ cells were observed. Looking at cell number within individual larvae in the same experiment as in Fig 3C (**Fig 3D**), we observe that at the given timepoints after amputation, eGFP^+^mCherry^+^ cells are the most abundant cell type at the wound site. The combined observations on cell motility, morphology and kinetics of recruitment at the wound site, suggest that non-motile eGFP^+^mCherry^-^cells with epithelial cell morphology are likely keratinocytes, whereas eGFP^+^mCherry^+^ are likely innate immune cells being recruited to the wound site. Thus, when considering motile cells, both transgenic lines mark similar innate immune cell populations, which increase their expression of *tnfb* after wounding. The *Tg(tnfb:eGFP-F)* line also marks a small subpopulation of epithelial cells, likely keratinocytes which, based on motility and morphology, can be distinguished from motile innate immune cells.

**Figure 3.**
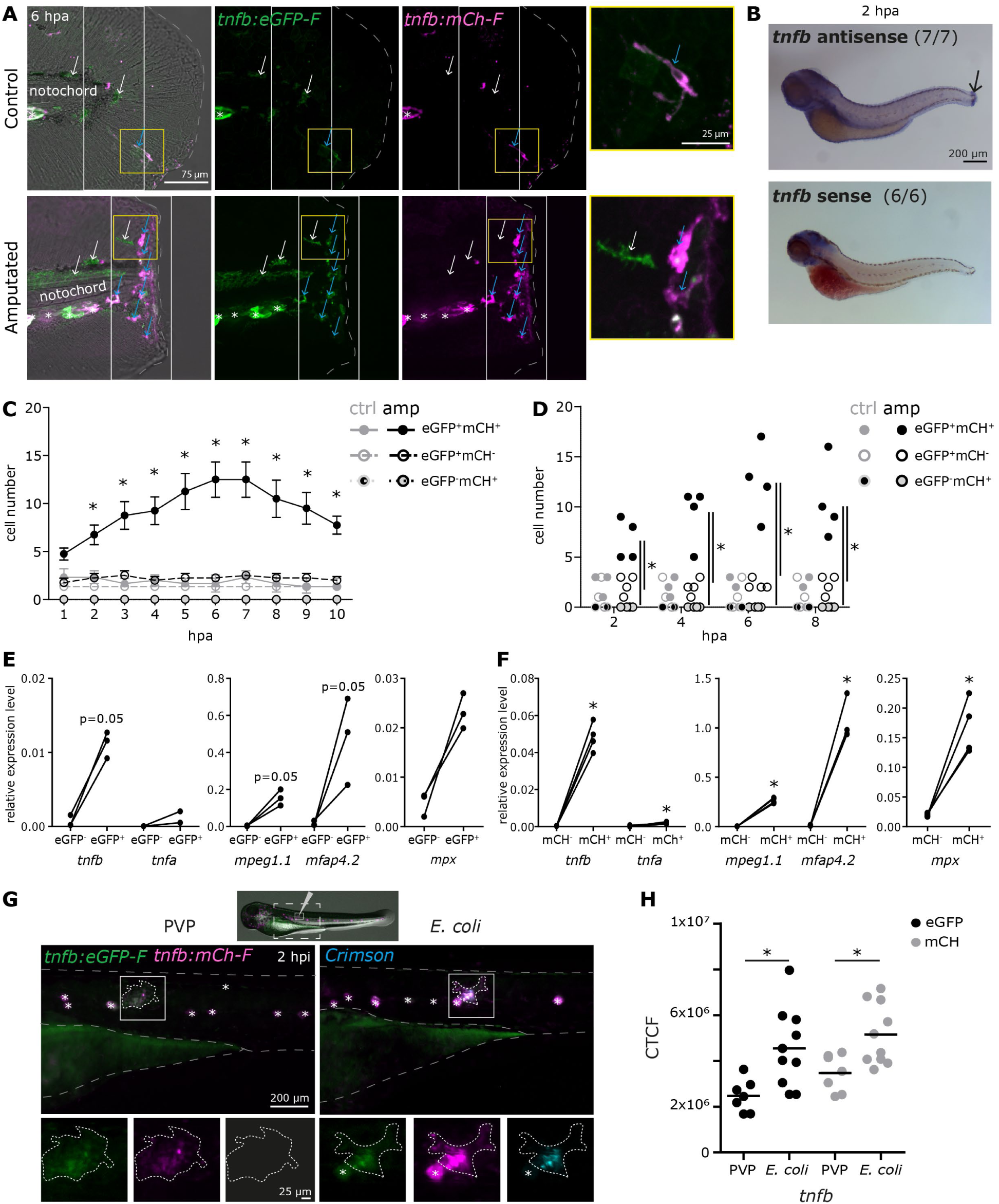
Reporter lines *Tg(tnfb:eGFP-F) and Tg(tnfb:mCherry-F)* recapitulate transcriptional activation of *tnfb* in innate immune cells after wounding. A-F) Larvae (2 dpf) were screened based on transgene expression in neuromasts. Caudal fins of 3 dpf larvae were amputated (amp) or left intact (control, ctrl). A) Confocal imaging of *Tg(tnfb:eGFP-F;tnfb:mCherry-F)*. The top panel shows control caudal fin with few fluorescent cells. Grey dashed line indicates the outer perimeter of the caudal fin. White square delimits the wound site used for cell counting, defined by a width of 105 µm centred around the tip of notochord. White arrows indicate eGFP^+^mCherry^-^cells, blue arrows indicate eGFP^+^mCherry^+^ cells clearly increasing in number after wounding (lower panel). Asterisks (*), indicate neuromasts. **B)** Representative images of zebrafish larvae acquired with Zeiss Stemi 508 stereo microscope after wholemount *in situ* hybridisation of *tnfb* antisense (top panel), *tnfb* sense probes (negative control, lower panel). *tnfb* expression was detected in the amputated fin (black arrow). **C)** Kinetics of *tnfb* expression were determined at the wound site by manually counting transgene-expressing cells. Graphs show mean and SEM. Asterisks (*) indicates significant differences, at a given time point, between the same cell type in the amputated and intact group, as assessed by Two-Way ANOVA followed by Tukey’s post-hoc test. **D)** Individual data derived from the same *tnfb* expression data as C), at the indicated timepoints, number of cells per cell type are shown. Asterisks (*) indicates significant differences within treatment at the indicated timepoint, as assessed by Two-Way ANOVA followed by Tukey’s post-hoc test. **E-F)** Relative expression level of *tnfb, tnfa, mpeg1.1, mfap4.2 and mpx* in **E)** eGFP^+^- or **F)** mCherry^+^-sorted fractions 2-3 hpa from *Tg(tnfb:eGFP-F)* or *Tg(tnfb:mCherry-F)* larvae, using *eef1a1l1* as a reference gene. Dots represent independent experiments; sorted fractions from the same experiment are connected. Asterisks (*) indicate significant differences as assessed by One-tailed Mann-Whitney test. **G)** *Tg(tnfb:eGFP-F;tnfb:mCherry-F)* larvae (3 dpf) were injected intramuscularly with 1 nl PVP or with 2×10^3^ CFU *E. coli* crimson resuspended in PVP. Images were acquired 2 hpi with a Leica M205FA fluorescence stereo microscope with 61x zoom. The area around the fluorescent reporter protein (dashed outline) was selected in FIJI for total fluorescent quantification. Asterisks (*), indicate neuromasts. **H)** Larvae were treated as in G. Corrected total cell fluorescence (CTCF) in PVP- or *E. coli*-injected individuals was measured within the dashed outlined area as described in G. Neuromasts were excluded from the total fluorescence quantification. Asterisks (*) indicate significant difference assessed by One-tailed Mann-Whitney test.

To further confirm that eGFP^+^ or mCherry^+^ cells express *tnfb*, prior to RT-qPCR (**Fig 3E-F**), sorting was performed to obtained eGFP^+^- and eGPF^-^- or mCherry^+^ and mCherry^-^cell fractions (**Supplementary Fig 1**) from pools of *Tg(tnfb:eGFP-F)* or *Tg(tnfb:mCherry-F)* zebrafish larvae (3 dpf) 2-3 hours post-amputation (hpa). eGFP^+^ or mCherry^+^ cells were enriched in *tnfb* transcripts and to a lesser extend in *tnfa* transcripts (**Fig 3E-F left panel**), confirming that in both transgenic lines *tnfb* expression is induced in reporter protein-positive cells early during inflammation. Next, to gain insights into the types of cells expressing *tnfb*, expression of typical macrophage (*mpeg1.1, mfap4.2*) and neutrophil (*mpx*) markers was assessed in the same sorted cells (**Fig. 3E-F**). eGFP^+^ or mCherry^+^ cells were not only enriched in *tnfb* transcripts, but also in *mpeg1.1, mfap4.2* (**Fig 3E-F middle panel**), and *mpx* transcripts (**Fig 3E-F right panel**). Altogether, these data suggest that, after wounding, macrophages and neutrophils are among the immune cells expressing *tnfb*. The specific expression of *tnfb* in innate immune cells agrees with data retrieved from the publicly available Daniocell dataset reporting on RNA expression in steady-state developing zebrafish ((Farrell et al., 2018; Sur et al., 2023), **Supplementary Fig 5**) and, within the dataset we additionally explored *tnfa* and *il1b* transcript expression. *tnfb* and *il1b* transcripts were expressed in subpopulations of neutrophils, macrophages, microglia and hematopoietic stem cells (myeloid), while *tnfa* transcripts were barely detectable, except in a few macrophages (**Supplementary Fig 5A**). In these *tnfa^+^* macrophages coexpression of *tnfb* or *il1b* transcripts occurred (**Supplementary Fig 5B left-middle panel**). The neutrophil population in general, was predominantly composed of *il1b^+^tnfb^-^*cells, with a small subpopulation *il1b^+^tnfb^+^* cells. In nearly all *tnfb^+^* neutrophils, macrophages, microglia and hematopoietic stem cell (myeloid), co-expression of *il1b* was observed (**Supplementary Fig 5B right panel**). Constitutive expression of *tnfb* could be detected in the lateral line cell cluster, however, the dataset did not facilitate distinction of mantle cells from other cell types within neuromast as separate populations (**Supplementary Fig 5C**). Prior to the emergence of any of the aforementioned immune cell types, *tnfb* was abundantly expressed in the periderm (**Supplementary Fig 5C**), suggesting that *tnfb* may play a role in early development. However, once innate immune cells are formed, they are the primary source of *tnfb*. Together, the data demonstrate that innate immune cells are the main cell sources of *tnfb* during development (Daniocell dataset) and in response to wounding (this study), thereby confirming that the motile *tnfb-*expressing cells recruited to the wound are indeed innate immune cells, and likely co-express *tnfa* and/or *il1b*.

To investigate whether *tnfb* expression in innate immune cells can be triggered not only after wounding but also after infection, intramuscular injection of *E. coli* was performed in 3 dpf larvae and PVP injection served as no bacteria control (**Fig 3G**). To quantify *tnfb:eGFP-F* and *tnfb:mCh-F* expression in innate immune cells recruited to the site of injection, the injection area was carefully selected (**Fig 3H**, area within dashed line) and neuromast were excluded from total fluorescent analysis. In both, PVP and *E. coli* injected larvae, *tnfb:eGFP-F* and *tnfb:mCh-F* were induced at 2 hours post injection (hpi), with the total eGFP and mCherry fluorescence being higher in the *E. coli* compared to the PVP injected group (**Fig 3H**). This data suggests that in the context of a bacterial infection, *tnfb* is transcriptionally induced during the early, proinflammatory phase, and that both reporter lines can be used to evaluate *tnfb* expression during infection. Altogether, during early zebrafish development innate immune cells constitutively express *tnfb*, and its expression can be induced in these cells after wounding or *E. coli* injection. When considering *tnfb* expression in (motile) immune cells, both transgenic lines recapitulate transcriptional activation of *tnfb.* The *Tg(tnfb:eGFP-F)* line also marks a population of non-immune cells, likely keratinocytes, which is not marked by the *Tg(tnfb:mCherry-F)* line, but can be distinguished from immune cells base on their morphology and motility. Combined, these transgenic lines are new tools, useful to study *tnfb* kinetics of expression and cell sources *in vivo* during inflammation.

### A subset of macrophages and of neutrophils express *tnfb* after wounding

The kinetics of *tnfb* expression were tracked in macrophages and neutrophils over a period of 6h after amputation, and the total number of macrophages, neutrophils and *tnfb*-expressing cells, was quantified at the 6h time point.

In intact caudal fins of *Tg(mpeg1:mCherry-F;mpx:eGFP*) larvae, macrophages (mCherry^+^) and neutrophils (eGFP^+^) were present in small number; after amputation, both significantly increased, with macrophages being generally more numerous than neutrophils (**Fig 4A-B**). When assessing *tnfb* expression in macrophages, in intact fins of *Tg(tnfb:mCherry-F;mpeg1:eGFP)* larvae, basal expression was observed in the majority of macrophages (**Fig 4C-D**); after amputation, the number of *tnfb:mCh-F-*expressing macrophages increased and a low number of *mpeg1:eGFP^-^tnfb:mCh-F^+^* cells (6.73% of *tnfb*^+^ cells) were also observed (**Fig 4C-D**). When looking at the kinetics of recruitment of *tnfb:mCh-F-*expressing macrophages (**Supplementary video 3**), *mpeg1:eGFP^+^tnfb:mCh-F* ^+^ cells were observed as early as 1 hpa, the earliest timepoint assessed, whereas *mpeg1:eGFP ^-^tnfb:mCh-F ^+^* cells were not observed until 4 hpa.

**Figure 4.**
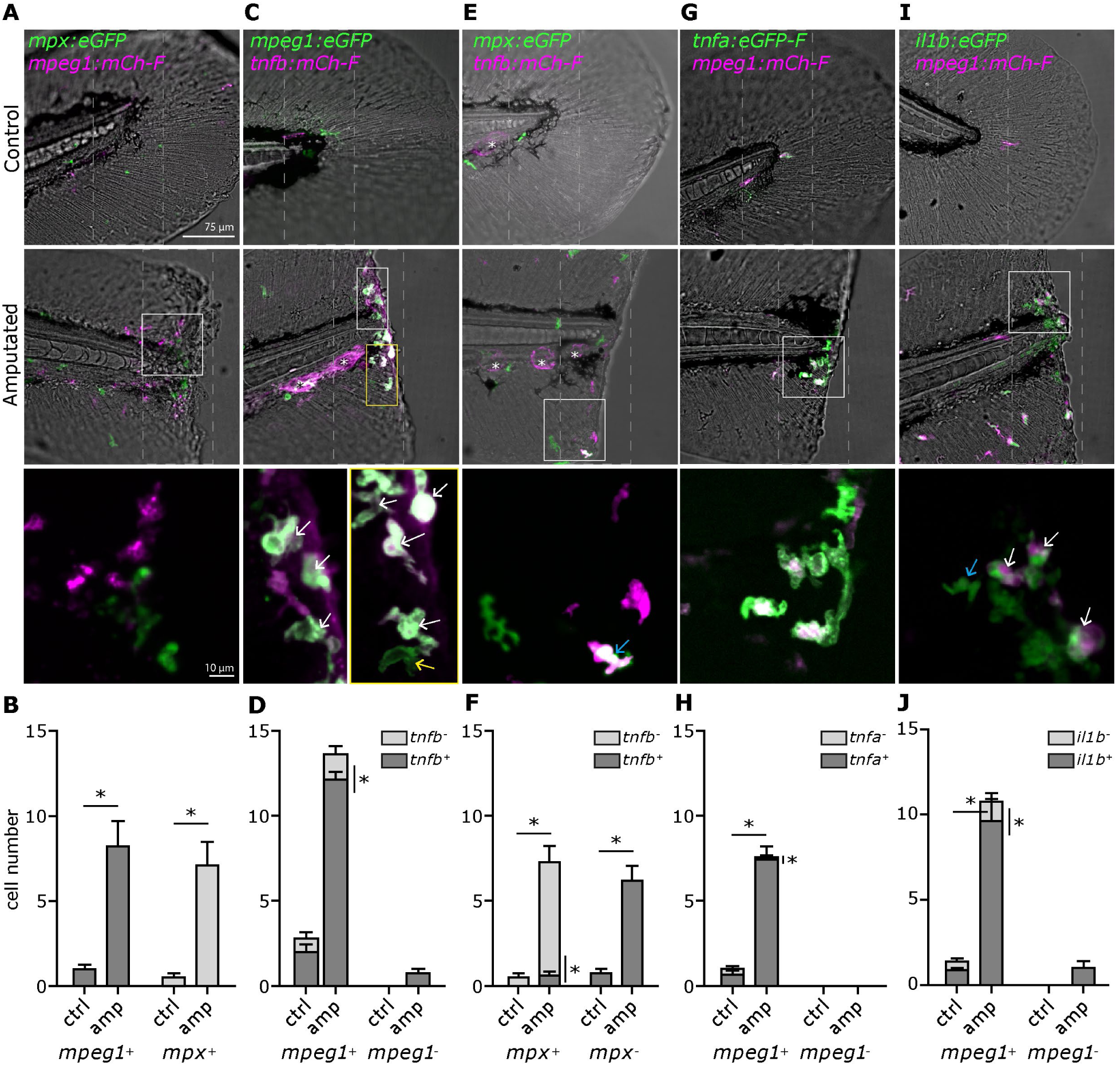
Macrophages and a subset of neutrophils express *tnfb:mCh-F* during wounding-induced inflammation. Caudal fins of 3 dpf transgenic larvae were amputated (amp) or left intact (control, ctrl), and imaged with an Andor spinning disk confocal microscope using 20x magnification (n=5-11 larvae per group). To aid cell counting, Z-stacks of 21 µm were acquired at a 6-minute interval between 5.5-6.5 hpa. In each pair of panels (**A-B** through **I-J**), the upper panel shows a representative image either of a control or of an amputated caudal fin (Asterisks (*) indicate neuromast); the lower bar graph panel, shows the corresponding total cell count. **A)** Macrophages (mCherry^+^) and neutrophils (eGFP^+^) are recruited to the wound and increase in number compared to the control. **C)** The majority of macrophages (eGFP^+^) express *tnfb* (eGFP^+^mCherry^+^; white arrows), while few eGFP^+^mCherry^-^(*mpeg1^+^tnfb^-^*) cells can be seen within the yellow outline. The yellow inset shows the same field of view marked by the yellow outline, 12 minutes later, with improved visibility of eGFP^+^mCherry^-^cells (yellow arrow). **E)** Only a subpopulation of neutrophils (eGFP^+^) express *tnfb* (eGFP^+^mCherry^+^; blue arrow) in response to wounding. **G)** Macrophages (mCherry^+^) were the only cells expressing *tnfa* (mCherry^+^eGFP^+^) **I)**. Macrophages (mCherry^+^) are the main producers of *il1b* (mCherry^+^eGFP^+^; white arrow), while few mCherry^-^eGFP^+^ (*mpeg1^-^il1b^+^*) cells can also be seen (blue arrow). **B/D/F/H/J)** Total cell count. Cells were counted manually within the grey dashed box. In **B**, asterisks (*) indicate significant differences between the control and amputated group within each cell type as assessed using Kluskal-Wallis test followed by a Dunn’s multiple comparison test; in **D/F/H/J**, significant differences were calculated within each cell type group, and between cell type groups within the same treatment as assessed using Kluskal-Wallis test followed by a Dunn’s multiple comparison test. Asterisks (*) indicates significant difference between the indicated groups.

When assessing *tnfb* expression in neutrophils, in intact fins of *Tg(tnfb:mCherry-F;mpx:eGFP)* larvae, none of the neutrophils were *tnfb:mCh-F ^+^* at the selected 6h timepoint, whereas after amputation, out of the total neutrophils recruited to the wound, a small subpopulation (8.75%) expressed *tnfb:mCh-F* (**Fig 4E-F**), suggesting that the small population of *mpeg1:eGFP ^-^tnfb:mCh-F ^+^* cells observed in **Fig 4C-D**, were likely neutrophils. When looking at the kinetics of *tnfb* expression in neutrophils, *mpx:eGFP^+^tnfb:mCh-F*^+^ cells were not observed at the wound site before 3 hpa (**Supplementary video 3**), showing that neutrophils express *tnfb:mCh-F* to a detectable level later than macrophages, as *mpeg1:eGFP^+^tnfb:mCh-F^+^* cells were observed from 1 hpa onwards. Together with our gene expression analysis (**Fig 3E-F**), these data show that after amputation both, macrophages and neutrophils, express *tnfb*, with macrophages being the main producers as *tnfb*-expressing macrophages were more numerous than *tnfb*-expressing neutrophils, and that *tnfb* expression was triggered earlier in macrophages than in neutrophils.

To compare cellular expression of *tnfb* to that of *tnfa* or *il1b*, we selected the 6 hpa timepoint, the time at which expression of all three transgenes is significantly increased after amputation. At this timepoint the total number of cytokine-expressing cells and the proportion of those that were macrophages, was quantified. To this end, we used double transgenic lines with macrophages in red and *il1b* or *tnfa* in green: *Tg(tnfa:eGFP-F;mpeg1:mCherry-F)* or *Tg(il1b:eGFP;mpeg1:mCherry-F).* When considering *tnfa* (**Fig 4G-H**), in the intact fin of *Tg(tnfa:eGFP-F;mpeg1:mCherry-F)* few macrophages were present, with the majority showing basal *tnfa:eGFP-F* expression. After amputation the number of recruited macrophages increased, with the majority being *tnfa:eGFP-F*^+^; only occasionally macrophages that were *tnfa:eGFP-F*^-^were observed (1.5%). *Mpeg1:mCh-F* ^-^*tnfa:eGFP-F*^+^ cells were not observed. When considering *il1b* (**Fig 4I-J**), in intact fins basal expression of *il1b:eGFP* was observed in *Tg(il1b:eGFP;mpeg1:mCherry-F).* After amputation the majority of recruited macrophages were *il1b:eGFP^+^* although a subpopulation of *mpeg1:mCh-F ^-^il1b:eGFP^+^* cells (12.7% of total *il1b^+^* cells) could be identified. Altogether, these data confirm that in response to injury *tnfb, tnfa,* and *il1b* are mainly expressed by macrophages. While *tnfa* is expressed exclusively by macrophages, *tnfb* and *il1b* are expressed by macrophages and also by a subpopulation of neutrophils.

### Kinetics of *tnfb, tnfa* and *il1b* expression after amputation

After having characterised which innate immune cells express *tnfb, tnfa* and *il1b,* and having shown cell-specific expression kinetics of *tnfb* (**Fig 4, Supplementary video 3**), we next characterised the kinetics of expression of all these transgenes and assessed whether they are expressed concomitantly or sequentially. To this end, crosses of lines with *tnfb-*expressing cells in fluorescent red and *tnfa-* or *il1b-*expressing cells in fluorescent green, *Tg(tnfa:eGFP-F);Tg(tnfb:mCherry-F)* or *Tg(il1b:eGFP);Tg(tnfb:mCherry-F)* were used. We imaged 3 dpf larvae over a period of 9h after amputation; to aid cell tracking and cell counting, images were acquired every 6 minutes. Owing to the availability of transgenic lines marking *tnfa* or *il1b* only in fluorescent green, it was not possible to assess co-expression of *tnfa* and *il1b*, or co-expression of all three cytokines within a cell.

When considering the kinetics of *tnfb* and *tnfa* expression (**Fig 5A and 5C**) in intact fins, only a few motile cells were observed and, if present, they were *tnfa:eGFP-F^-^tnfb:mCh-F^+^.* In amputated fins, the majority of transgene-expressing motile cells were *tnfa:eGFP-F^+^tnfb:mCh-F ^+^* and from 3 hpa onward their number significantly increased compared to the same cells type in the intact fins. A population of *tnfa:eGFP-F^-^tnfb:mCh-F^+^* cells was also identified, most prevalent at 2 hpa. When these cells were tracked (**Supplementary Video 4**), we observed that many transitioned into *tnfa:eGFP-F^+^tnfb:mCh-F^+^* cells. This pattern of cytokine expression, combined with our observation that *tnfa:eGFP-F* is exclusively expressed by macrophages (**Fig 4G-H**), suggests that these are macrophages in which *tnfb* is expressed prior to *tnfa.* Conversely, *tnfa:eGFP-F^-^tnfb:mCh-F^+^* cells present after 4 hpa remained *tnfa:eGFP-F^-^tnfb:mCh-F^+^*. *tnfa:eGFP-F^+^tnfb:mCh-F^-^*cells were not observed.

**Figure 5.**
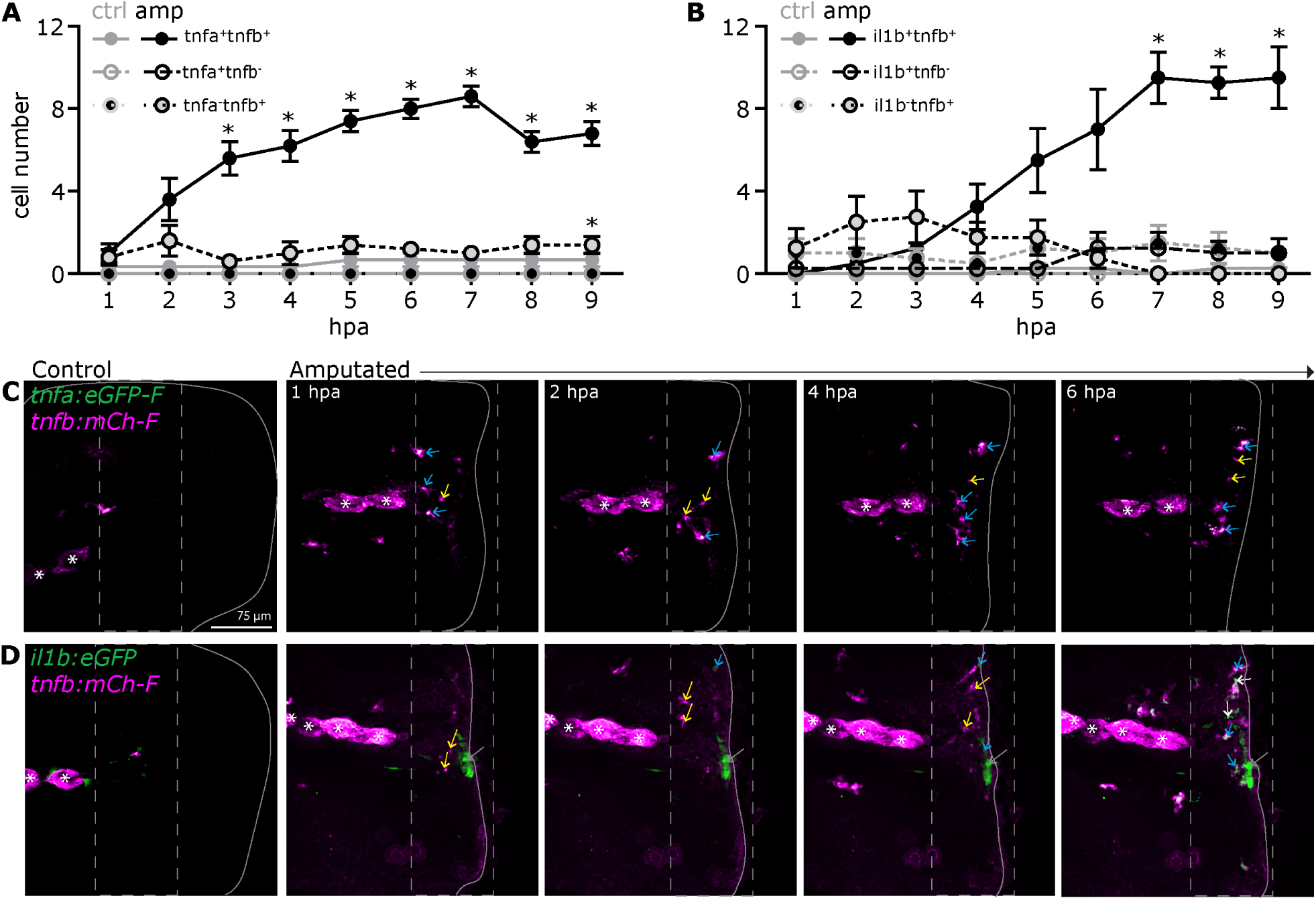
Kinetics of *tnfb, tnfa* and *il1b* expression in innate immune cells after amputation. Caudal fins of 3 dpf transgenic larvae were amputated (amp) or left intact (control, ctrl) and imaged during 9 hpa with an Andor spinning disk confocal microscope using 20x magnification (n=4-5 larvae per group). To aid manual counting of cells, Z-stacks of 21 µm were acquired at a 6-minute interval. **A-B)** Kinetics of expression of the indicated cytokines were determined at the wound site by manually counting transgene-expressing cells. Graphs show mean and SEM. Asterix (*) indicates significant differences between the same cell type in amputated and control fins at the same timepoint, as assessed by 2-WAY ANOVA with Geisser-Greenhouse correction for sphericity, followed by Tukey’s multiple comparisons test. **C**) Representative maximum projections of a control or amputated (grey outline) fin of *Tg(tnfa:eGFP-F);Tg(tnfb:mCherry-F)* larvae. Left panel shows a control fin; panels on the right show an amputated fin of the same individual at the indicated timepoints. Arrows indicate typical *tnfa:eGFP-F^+^tnfb:mCh-F^+^* (cyan arrow) and *tnfa:eGFP-F^-^tnfb:mCh-F^+^* (yellow arrow) cells. **D**) *Tg(il1b:eGFP);Tg(tnfb:mCherry-F)* larvae treated as in C. Arrows indicate typical *il1b:eGFP^+^tnfb:mCh-F^+^* (cyan arrow), *il1b:eGFP^-^tnfb:mCh-F^+^* (yellow arrows), and *il1b:eGFP^+^tnfb:mCh-F^-^*(white arrow) or immotile *il1b:eGFP^+^tnfb:mCh-F ^-^*(grey arrow) cells.

When considering the kinetics of *tnfb* and *il1b* expression (**Fig 5B and 5D**) in intact fins, only a few transgene-expressing motile cells were observed, the majority of which were *il1b:eGFP^-^tnfb:mCh-F ^+^*, and a few were *il1b:eGFP^+^tnfb:mCh-F ^+^*. *il1b:eGFP^+^tnfb:mCh-F ^-^*cells were not observed. After amputation, between 1-3 hpa, the majority of cells present at the wound were *il1b:eGFP^-^tnfb:mCh-F^+^* cells. Many of these cells transitioned into *il1b:eGFP^+^tnfb:mCh-F ^+^* cells (**Supplementary Video 4**), the main population present by 4 hpa onwards. From 6 hpa onwards, a small population of *il1b:eGFP^+^tnfb:mCh-F^-^*cells is present at the wound. Together with macrophages as the main cell type expressing *il1b:eGFP* (**Fig 4J**), this suggests that the largest population, *il1b:eGFP^+^tnfb:mCh-F^+^* cells, is mostly composed of macrophages in which *tnfb* expression is detected earlier than *il1b* expression. Altogether, this suggests that the temporal dynamic of cytokine expression in macrophages is sequential, with of *tnfb* first, followed by *tnfa* and *il1b*.

### *tnfb* expression by innate immune cells following *E. coli* injection

After having established that *tnfb:eGFP-F* and *tnfb:mCherry-F* are induced during the early phase of *E. coli*-induced inflammation (**Fig 3G-H**), we investigated whether *tnfb* is expressed with *tnfa* and *il1b.* To this end, transgenic larvae (3 dpf) were injected intramuscularly with *E. coli* or with PVP (control) and imaged at 6 hours post injection (hpi). Since at this stage it is challenging to manually count cells due to recruited cells crowding the injection area total fluorescence quantification instead of single cell count was performed. We made use of *Tg(tnfa:eGFP-F);Tg(tnfb:mCherry-F)* or *Tg(il1b:eGFP);Tg(tnfb:mCherry-F)* larvae. Based on total fluorescence of *tnfa:eGFP-F, il1b:eGFP* as well as *tnfb:mCh-F,* their expression increased significantly in the *E. coli*-injected compared to their corresponding PVP-injected group (**Fig 6A-D**). These data indicate that, like *tnfa* and *il1b, tnfb* is also induced following *E. coli* injection.

**Figure 6.**
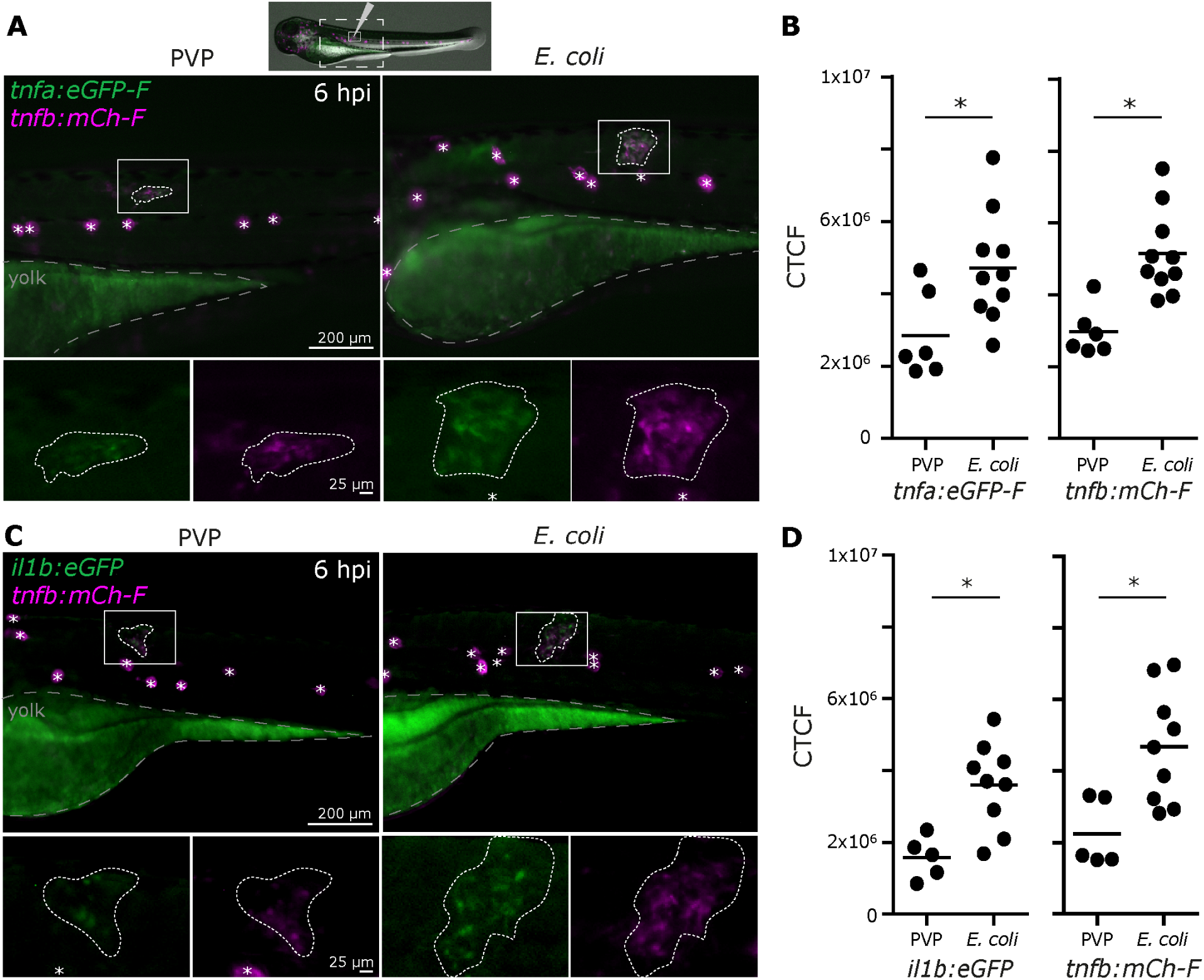
Expression of *tnfb, tnfa* and *il1b* after E. coli injection. Transgenic zebrafish (3 dpf) were injected intramuscularly with 1 nl PVP or with 2×10^3^ CFU *E. coli* crimson resuspended in PVP, and images were acquired at 6 hpi with a Leica M205FA fluorescence stereo microscope using 58-61x digital zoom. In each pair of panels **A-B)** *Tg(tnfa:eGFP-F);Tg(tnfb:mCherry-F)* or **C-D**) *Tg(il1b:eGFP);Tg(tnfb:mCherry-F)*, the left panel (**A/C**) shows representative images of PVP- or *E. coli*-injected larvae (Asterisks (*) indicate neuromast); the right panel (**B/D**) shows the corresponding total green or red fluorescence for the indicated cytokine-reporter measured within the dashed outlined area. Neuromasts were excluded from total fluorescence quantification. Asterisks (*) indicate significant difference assessed by one-tailed Mann-Whitney test.

Next, we assessed which innate immune cells expressed *tnfb:mCh-F, tnfa:eGFP-F,* or *il1b:eGFP* at 2.5 hours post injection with PVP or *E. coli*. In *Tg(mpeg1:mCherry-F;mpx:eGFP*) larvae, both macrophages (mCherry^+^) and neutrophils (eGFP^+^) were observed at the injection site (**Fig 7A**). When assessing *tnfb* expression, after PVP injection, *tnfb:mCh-F* expression was observed in macrophages (*mpeg1:mCh-F^+^*) (**Fig 7B, Supplementary video 5**). When using the *Tg(tnfb:mCherry-F;mpx:eGFP)* line, no neutrophils were observed which express *tnfb:mCh-F* after PVP injection, whereas *tnfb:mCh-F^+^* neutrophils could be observed after *E. coli* injection (**Fig 7C**). When assessing *tnfa* expression by macrophages, in both injected groups, *mpeg1:mCh-F^+^tnfa:eGFP-F^+^* cells were observed (**Fig 7D, Supplementary video 5**). When assessing *il1b* expression by macrophages, *mpeg1:mCh-F^+^il1b:eGFP^+^* were observed after PVP injection, whilst after *E. coli* injection both *mpeg1:mCh-F^+^il1b:eGFP^+^* and *mpeg1:mCh-F ^-^il1b:eGFP^+^* cells were observed (**Fig 7E**). This suggest that in response to PVP or *E.* coli, macrophages are the cells expressing *tnfb, tnfa* and *il1b.* In addition to macrophages, a subset of neutrophils expresses *tnfb* after *E. coli* injection.

**Figure 7.**
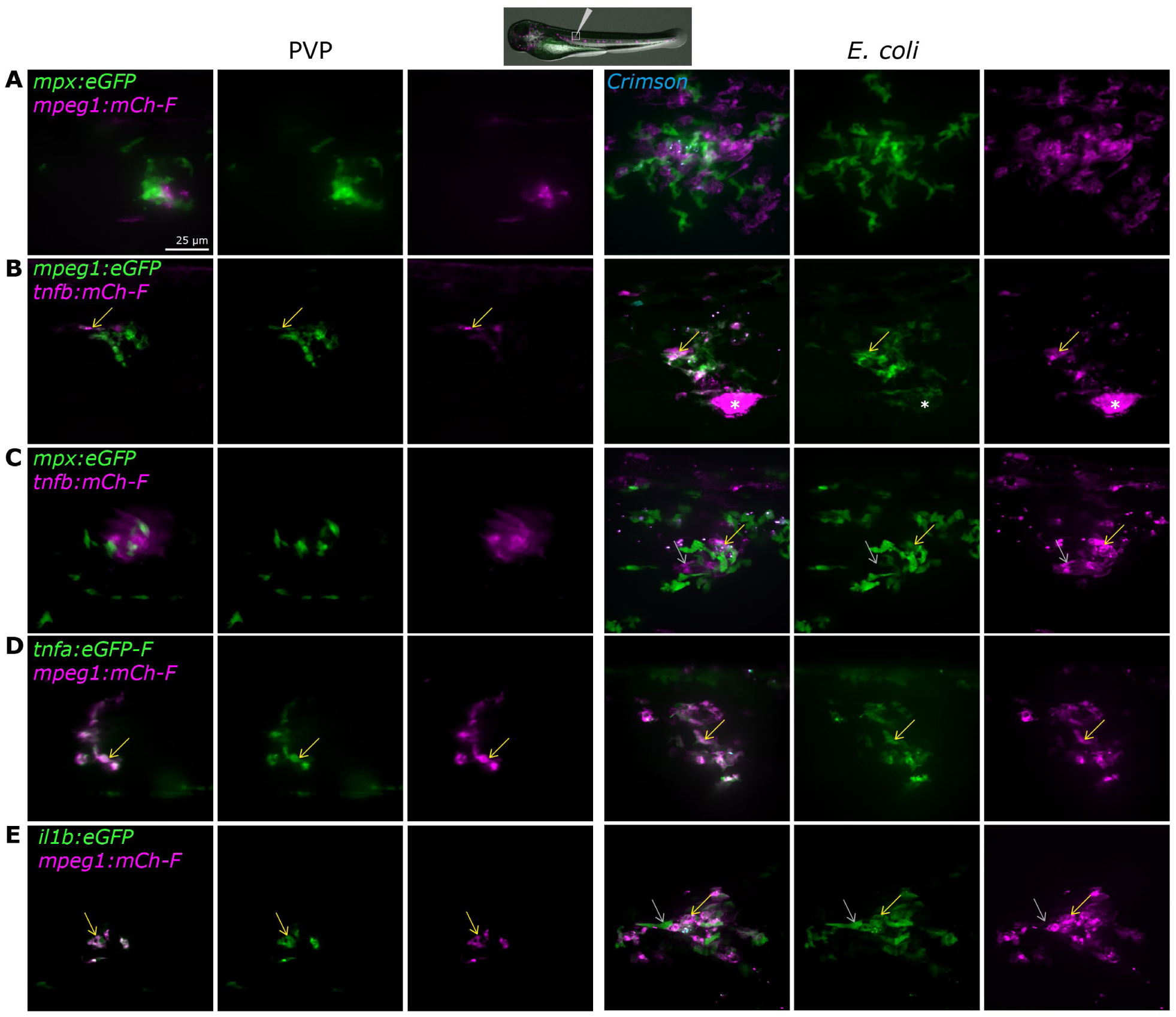
Occurrence of innate immune cells expressing *tnfb, tnfa* and *il1b* after *E. coli* injection. Transgenic zebrafish (3 dpf) were injected intramuscularly with 1 nl PVP or with 2×10^3^ CFU *E. coli* crimson resuspended in PVP, and imaged at 2.5 hpi with an Andor spinning disk confocal microscope using 40x (A, C-E) or 20x (B) magnification. Representative maximum projections of PVP- and *E. coli*-injected larvae, using Z-stacks of 25-40 µm are shown. **A)** Macrophages (mCherry^+^) and neutrophils (eGFP^+^) are observed after PVP injection and in response to *E. coli*. **B)** Macrophages (eGFP^+^) express *tnfb* (eGFP^+^mCherry^+^; yellow arrow) and a second *tnfb-expressing* cell population is observed (eGFP^-^mCherry^+^; grey arrow) (Asterisks (*) indicate neuromast). **C)** Only a subpopulation of neutrophils (eGFP^+^) express *tnfb* (eGFP^+^mCherry^+^; yellow arrow) in response to *E. coli.* **D)** Macrophages (mCherry^+^) express *tnfa* (mCherry^+^eGFP^+^; yellow arrow), and **E)** Macrophages (mCherry^+^) express *il1b* (mCherry^+^eGFP^+^; yellow arrow), and mCherry^-^eGFP^+^ (*mpeg1^-^il1b^+^*; grey arrow) cells are observed.

Combined, these data are consistent with and further extend our findings that during the early phase of inflammation *tnfb* is induced and modulated *in vivo*, in subpopulations of macrophages and neutrophils.

## Discussion

Despite zebrafish possessing two homologues of human *TNF*, *tnfa* and *tnfb* (Kinoshita et al., 2014), research has largely focussed on *tnfa,* leaving *tnfb* underexplored. In this study, we generated two *tnfb* reporter lines and used them to identify *tnfb*-expressing cells and to describe *tnfb* expression kinetics *in vivo* during inflammation. We report on the constitutive *tnfb* expression in neuromast mantle cells, and inducible *tnfb* expression in innate immune cells during inflammation induced by caudal fin fold amputation and *E. coli* injection. By combining our newly generated *tnfb* reporters with available *tnfa* or *il1b* reporter lines and high-resolution microscopy, we report that macrophages are the main cells expressing *tnfa, tnfb, and il1b*, while neutrophils can express *tnfb* and *il1b,* but did not *tnfa* in response to wounding or to bacterial injection. We also show that the timing of *tnfb* expression differs between macrophages and neutrophils. Together, these results reveal cell type-specific expression of *tnfb* and distinct expression kinetics of *tnfa* and *tnfb,* setting the stage for future studies in which both homologs should be investigated. This will be essential to fully capture their overlapping as well as distinct roles during inflammation or infection, and how these relate to the better-known pleiotropic nature of mammalian TNF.

Here, we developed two transgenic reporter zebrafish lines (*Tg(tnfb:eGFP-F)* and *Tg(tnfb:mCherry-F)*); one, in which a 3.2-kb portion of the predicted *tnfb* promotor drives farnesylated eGFP (eGFP-F) expression and a second in which a 6.2-kb portion of the predicted *tnfb* promotor drives farnesylated mCherry (mCherry-F) expression. Direct comparison of fluorescent cells in crosses of these novel reporter lines, showed that both lines report constitutive expression of *tnfb* in neuromast mantle cells and inducible transcription of *tnfb* in innate immune cells during inflammation. Likely due to missing regulatory elements in the shorter promotor or an insertional effect, the *Tg(tnfb:eGFP-F)* line also marks a small subpopulation of keratinocyte-like cells that, based on motility and morphology, are easily distinguishable from motile innate immune cells. Nevertheless, *eGFP-F* and *mCh-F* expression kinetics were similar in immune cells. Together, *Tg(tnfb:eGFP-F)* and *Tg(tnfb:mCherry-F)* are novel complementary tools for characterising similarities and differences between *tnfa* and *tnfb* expression, in a range of cell types and inflammatory contexts.

By using two well-established inflammation models, we were able to contextualize *tnfb* expression and the immune cells involved. When focusing on macrophages in larval zebrafish, we report co-expression of *tnfb, tnfa* and *il1b* during the early phase of inflammation. These data agree with several previous studies reporting expression of the same cytokines in *tnfa:eGFP-F^+^* sorted macrophages 6h after amputation (Nguyen-Chi et al., 2015), in RNA sequencing (RNAseq) data of pools of sorted macrophages after *M. marinum* infection (Rougeot et al., 2019), in single-cell RNA-seq (scRNAseq) data of macrophages 3h after amputation (García-López et al., 2023), and in gene expression and RNAseq data of FACS-sorted macrophages 2h after amputation (Begon-Pescia et al., 2026). These studies consistently point towards *tnfb* also being expressed by macrophages, but did not study cytokine expression kinetics during inflammation, as they analysed a limited (2-3) number of timepoints, and at each timepoint, individuals were pooled. In this study, leveraging the availability of *tnfa:eGFP-F* and *il1b:eGFP* transgenic lines, we show that *tnfb:mCh-F^+^* cells can transition to *tnfa:eGFP-F-* or *il1b:eGFP-*expressing cells. Owing to the availability of transgenic lines marking *tnfa* or *il1b* only in fluorescent green, it was not possible to assess co-expression of *tnfa* and *il1b*, or co-expression of all three cytokines within a cell. However, our observation of distinct expression kinetics of the aforementioned cytokines combined with our observation that macrophages are the main *tnfb-expressing* cells after amputation, show that the temporal dynamics of the cytokines of interest, at least in macrophages, are sequential with expression of *tnfb,* followed by that of *tnfa* and *il1b*.

While we observed that *tnfa:eGFP-F* was expressed exclusively by macrophages, *tnfb:mCh-F* and *il1b:eGFP* were expressed by macrophages and by other cells. Others demonstrated that neutrophils indeed express *il1b* after amputation and during *Mycobacterium marinum* infection (Ogryzko et al., 2019). To our knowledge, this is the first report showing *in vivo* expression of *tnfb* by neutrophils and opens the possibility to use *tnfb* as a marker of neutrophil heterogeneity during inflammation. In mammals, it is increasingly appreciated that neutrophils occur with different activation states (Grieshaber-Bouyer et al., 2021; Rosales, 2018), among which a proinflammatory *TNF*-expressing subset has been observed (Khoyratty et al., 2021). *tnfb* expression as potential marker of neutrophils activation in response to amputation is reinforced by a study in larval zebrafish showing that, based on a photo-conversion lineage-tracing approach combined with single-cell RNAseq analysis, both rostral blood island- and caudal hematopoietic tissue-derived neutrophils are recruited to the wound and are enriched in *tnfb* (García-López et al., 2023). This suggests that *tnfb* expression is wound context-specific and independent of ontogeny. Furthermore, enrichment of *tnfb* in wound-responsive neutrophils, is conserved across developmental stages as demonstrated by heart-wound responsive neutrophils of adult zebrafish being enriched in *tnfb* (Wei et al., 2023). Based on co-expression with *il1b*, phagosome-related genes *ncf1* and *ncf2*, and chemotaxis genes *cxcr1, atf3 and illr4,* it was suggested that these subsets of neutrophils promote inflammation by increasing recruitment and retention of neutrophils at the wound (Wei et al., 2023). Together, this supports that *tnfb* expression reflects a proinflammatory neutrophil activation state in larval and adult zebrafish and, when used as a marker, may provide novel insights into neutrophils heterogeneity during inflammation.

We observed that *tnfb:mCh-F* expression was detected earlier in macrophages than in neutrophils involved in the same inflammatory response. Cell-specific kinetics of *tnfb* expression may indicate that *tnfb* expressed by macrophages has a role in inflammation initiation, while *tnfb* expressed by neutrophils has a role during inflammation amplification. In mammals, TNF signalling occurs via TNFRSF1A (previously TNFR1) or TNFRSF1B (previously TNFR2) (Horiuchi et al., 2010). TNFRSF1A is characterized by an intracellular death domain, which eventually leads to proinflammatory or apoptotic signals, while TNFRSF1B does not harbour an intracellular death domain and instead leads to the induction of proinflammatory and survival signals (Holbrook et al., 2019; Preedy et al., 2024; Zelová and Hošek, 2013). Zebrafish also possess Tnfrsf1a and Tnfrsf1b. Similar to mammalian TNFRSF1 expression patterns, also in zebrafish, *tnfrsf1a* is broadly expressed across several cell types and tissues, whereas *tnfrsf1b* has limited expression, mostly restricted to immune cells (Craig Smith and Farmh, 1994). These expression patterns are consistent with those retrieved from publicly available RNAseq datasets of steady-state developing zebrafish larvae (Farrell et al., 2018; Lange et al., 2024; Sur et al., 2023). Although it is likely that distinct Tnf-Tnfr interactions contribute to the pleiotropic downstream effects attributed to Tnf, ligand-receptor preferences or how receptor expression is modulated on different cell types during inflammation, remain to be elucidated.

Besides innate immune cells, we (this study) and others (Baek et al., 2022; Lush et al., 2019) found that neuromast mantle cells constitutively express *tnfb.* Based on hair cell ablation experiments, it was shown that in mantle cells *tnfb* can be further induced in response to wounding (Baek et al., 2022). Combined, this suggests a role for *tnfb* in neuromast maintenance and/or a protective role during the response against environmental stimuli. From a neuromast maintenance perspective, Tnfb could be involved in directing hair cell polarity. Zebrafish modified to constitutively express Tnfrsf1b show asymmetry between rostrad- and caudad-oriented hair cells, whereas these are symmetrically distributed in steady-state neuromasts (Jacobo et al., 2019). Another possibility is that mantle cells are thought to serve as stem cells to various cell types within neuromast. In fact, a recent study has shown that maintenance of their stem cell identity is dependent on another TNF superfamily member Tnfsf10 (previously Trail) (Fan et al., 2025), but whether Tnfb (Tnfs2) plays a role in stem cell maintenance remains to be established.

Examination of *tnfb* expression in publicly available scRNAseq datasets of steady-state developing zebrafish (Farrell et al., 2018; Sur et al., 2023) revealed its expression in periderm prior to immune cell development. Initially, *tnfb* expression was suggested to be a hallmark during the gastrulation phase of zebrafish development (Ito et al., 2008). This was plausible in light of mammalian TNF playing a role during organogenesis (You et al., 2021). However, others also found that *tnfb* expression during this early developmental timeframe is derived from periderm rather than the inner cell mass (Hoijman et al., 2021) suggesting a non-developmental role for *tnfb*. Periderm, otherwise known as the outermost protective layer of developing larvae (Liu et al., 2020), constitutively expresses *tnfb* at least until 5 dpf (Farrell et al., 2018; Hoijman et al., 2021; Sur et al., 2023). The widespread constitutive expression of *tnfb* in periderm, in mantle cells and in cells lining the exterior of the developing larvae, suggests a putative role for Tnfb in the response to stimuli from the aquatic environment.

In addition to constitutive *tnfb* expression, transient upregulation of *tnfb,* but not *tnfa,* was shown in mantle cells after hair cell ablation (Baek et al., 2022), again suggesting that Tnfb in particular may be involved in the response to environmental stimuli or wounding. When considering bacterial challenges, others have shown that *tnfb* was triggers upon both, immersion and injection of 3 dpf larvae with *Vibrio parahaemolyticus,* while *tnfa* was only enriched in injected larvae (Guo et al., 2021). It is possible that after immersion and injection different cell types are responsible for *tnfb* expression, with cells lining the exterior of the fish playing a main role after immersion and innate immune cells playing a role after injection.

Interestingly, besides zebrafish (cyprinid fish), differential expression of group I and group II Tnf may also occur in meagre, belonging to perciform fish. Like zebrafish, meagre also possess two TNF homologs called Tnf1 (group I) and Tnf2 (group II) (Milne et al., 2017). *In vitro* stimulation of primary gill cultures and intestinal cell suspension with several pathogen-associated molecular patterns (PAMPs) elicited increased expression biased towards Tnf1 (group I) rather than Tnf2 (group II). In contrast, stimulation of primary head kidney cells or splenocytes elicited a relatively large increase in expression for Tnf2 (Milne et al., 2017). Similar results were observed in other teleost species (Cui et al., 2020; Hong et al., 2013; Kadowaki et al., 2009; Kajungiro et al., 2015; Pleić et al., 2015; Zou et al., 2002). Altogether, this not only confirms that group I and II Tnf in fish are differentially regulated but is also indicative of a possible functional diversification between the two Tnf groups.

Both mammals and teleost possess complex repertoires of TNFSF ligands and their corresponding receptors (Goetz et al., 2004; Y. Li et al., 2021; Secombes et al., 2016). As a consequence of the teleost-specific whole-genome duplication event however, two or multiple copies of mammalian homologues may exist in fish. Duplicated genes may undergo various evolutionary fates: non-functionalisation (loss-of-function), neofunctionalisation (new function), or subfunctionalisation (Mihajlovic et al., 2025; OHNO et al., 1968). Subfunctionalisation is defined as the ancestral gene expression pattern or domain functions being divided between the duplicates (Lynch and Conery, 2000). The cell-specific and time-dependent expression pattern of *tnfa* and *tnfb* (this study) suggest a subfunctionalisation of gene expression. Whether this also has functional consequences, will be of interest going forward. Several examples of other zebrafish genes having undergone subfunctionalisation have recently been reviewed by *Tasnim et al*. Notably, divergence in spatial and temporal expression patterns have been leveraged to resolve aspects of mammalian protein function, structure and regulation (Tasnim et al., 2024), which would have been difficult, if not impossible, when such roles were combined into a single molecule.

In conclusion, we report on the constitutive *tnfb* expression in neuromast mantle cells, and on the inducible *tnfb* expression in subpopulations of macrophages and of neutrophils after wound- and bacterial-induced inflammation. *tnfb* kinetics of expression are cell-dependent, with expression in macrophages occurring prior to that in neutrophils. We also show that, at least in macrophages, *tnfb:mCh-F, tnfa:eGFP-F,* and *il1b:eGFP* are expressed sequentially, indicative of a differential role of these cytokines. Altogether, we propose that with the addition of *Tg(tnfb:eGFP-F)* and *Tg(tnfb:mCherry-F)* reporter lines, the zebrafish model is especially suited to dissect the pleiotropic nature of Tnf during infection or inflammation.

## Acknowledgments

This work was supported by the WIAS PhD grant from Wageningen University and by the European Union’s Horizon 2020 research and innovation programme through the Marie-Curie Innovative Training Network INFLANET (955576). The authors like to thank the CARUS Aquatic Research Facility of Wageningen University for fish rearing and husbandry, and Wageningen Light Microscopy Centre and Microspectroscopy Research Facility, and the Aquatic model facility ZEFIX from LPHI, University of Montpellier and the imaging facility BioCampus Montpellier Ressources Imagerie (MRI), member of the national infrastructure France-BioImaging supported by the French National Research Agency (ANR-10-INSB-04, “Investments for the future”).

## Supplementary figures

**Supplementary Figure 1.**
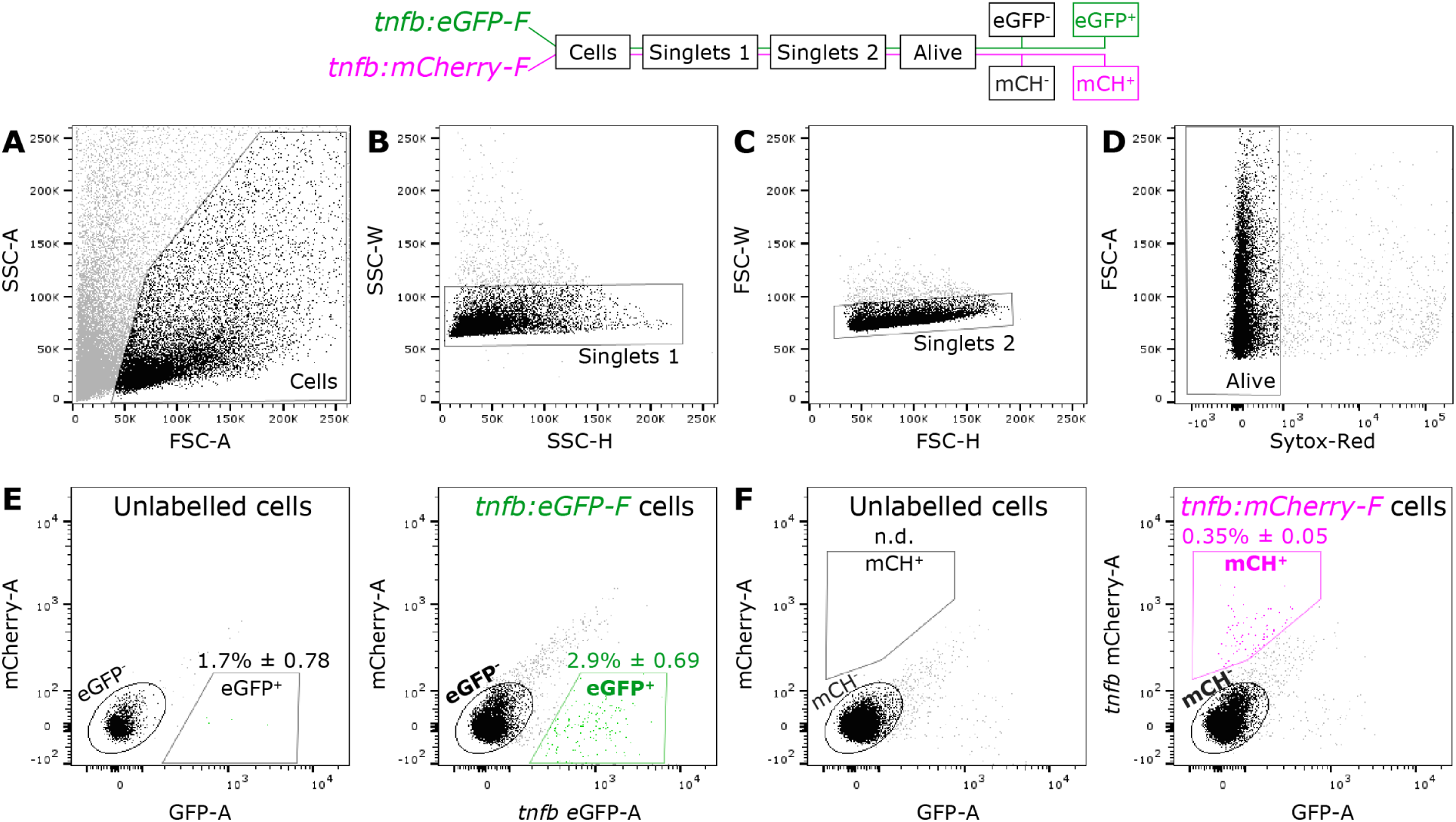
Fluorescence-Activated Cell Sorting (FACS) gating strategy to identify *tnfb-expressing* cells. Caudal fins of *Tg(tnfb:eGFP-F) or Tg(tnfb:mCherry-F)* larvae (3 dpf) were amputated, and 2-3 hpa FACS was used to sort eGFP^+^ and eGFP^-^fractions, or mCherry^+^ and mCherry^-^fractions, from whole larvae (n=300). Representative dot plots showing the gating strategy: **A**) Debris and dead cells were excluded based on the Forward (FSC-A) and Side Scatter (SSC-A) Area. **B**) Single cells were discriminated from doublets and clumps based on SSC-Width (SSC-W) vs SSC-Height (SSC-H) corresponding to Singlets 1 and **C**) by plotting FSC-Width (FSC-W) vs FSC-Height (FSC-H) corresponding to Singlets 2. **D**) Live cells were discriminated from dead cells based on Sytox Red viability staining, and Sytox Red–positive cells were excluded. **E**) Cells from wildtype larvae (Unlabelled cells (left panel)), used to define gating boundaries for sorted eGFP^+^ and eGFP^-^fractions (right panel). Percentages (%) represent mean ± SEM of 3 independent experiments. **F**) Cells from wildtype larvae (Unlabelled cells (left panel)), used to define gating boundaries for sorted mCherry^+^ and mCherry^-^fractions (right panel). Percentages (%) represent mean ± SEM of 4 independent experiments; n.d. = non-detectable. To be more stringent on purity of sorted populations, gates have been designed to not overlap. Gene expression analysis for sorted populations is shown in main Figure 3E-F.

**Supplementary Figure 2.**
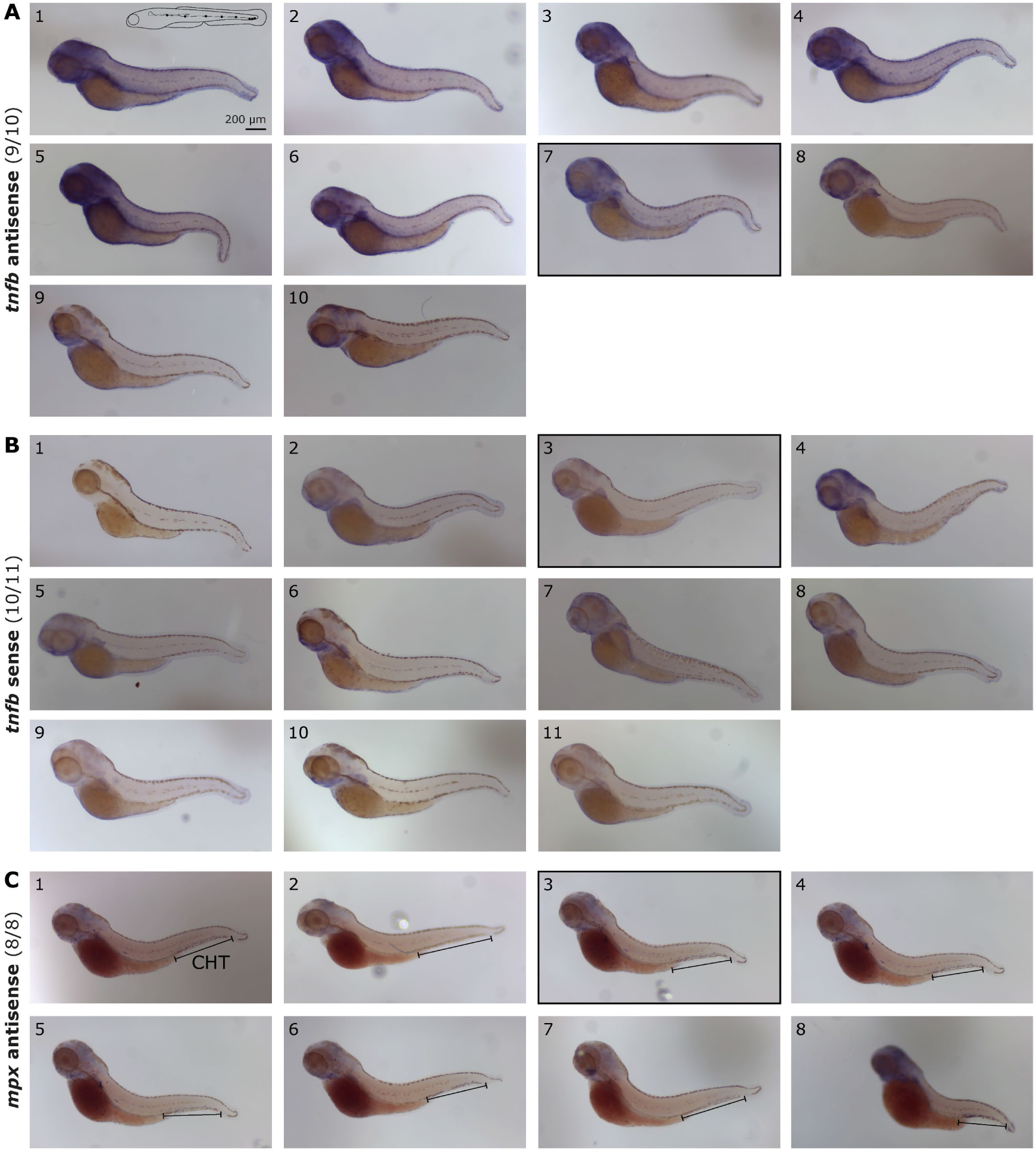
Wholemount *in situ* hybridization of *mpx* and *tnfb* mRNA in 3 dpf zebrafish. **A)** Panel of *tnfb* mRNA expression. Larvae were scored positive when signal was visible in at least two neuromast of the posterior lateral line. Positions of lateral neuromasts in 3dpf zebrafish are depicted in the schematic drawing. 9 out of 10 larvae were positive, only A9 did not meet this criterium. **B)** *tnfb sense* probe did not show any specific signal in the neuromast. B1 did not show any coloration, different from the remaining larvae where some coloration in the head was observed. **C)** Panel of *mpx* mRNA expression. Larvae were scored positive when signal was visible in the caudal hematopoietic tissue (CHT). All larvae (8/8) met this criterium. Black outlined are the images of larvae also shown in main Figure 1D.

**Supplementary Figure 3.**
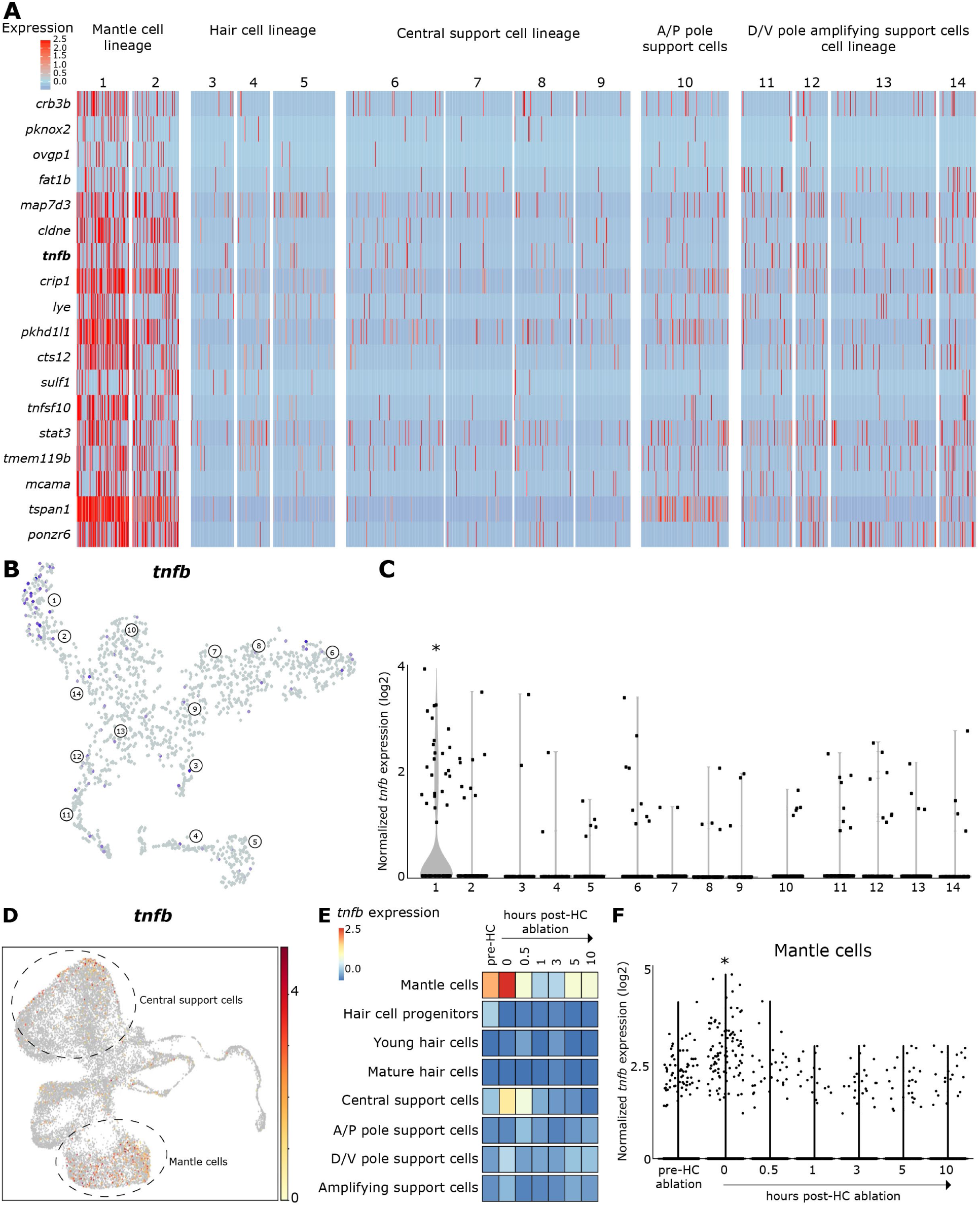
Mantle cells in homeostatic neuromasts or soon after hair cell ablation are enriched in *tnfb.* Single-cell RNA-sequencing data of homeostatic neuromasts (**A-C**) was retrieved from <u>Zebrafish Neuromast scRNAseq</u>, and data from neuromasts at multiple timepoints after hair cell (HC) ablation (**D-E**) was retrieved from <u>Neuromast regeneration scRNAseq</u>. **A)** Heatmap of mantle cell’s lineage markers, including *tnfb.* Heat bar shows log2 fold change of transcripts significantly (p<0.001) enriched in mantle cells as reported in Lush et al. 2019. Numbers 1-14 indicate cell clusters. Clusters under the same lineage represent a different developmental stage, for example clusters 3-5 are progenitor, young and mature hair cells, respectively. For the other lineages, this distinction is not as clear, including clusters 1-2, both marking the development of mantle cells. A/P: anterior-posterior, D/V: dorsal-ventral. **B)** t-distributed Stochastic Neighbour Embedding (t-SNE) plot of *tnfb*-expressing cells in blue, which are mostly located in the mantle cell’s lineage clusters 1 and 2. **C)** Violin plot of normalised *tnfb* expression per cell cluster. Mantle cell cluster 1 contains the highest number of *tnfb-*expressing cells and is significantly enriched with this gene. Asterisks (*) indicate significant difference (p<0.001) as assessed by Wilcoxon Rank test followed by a Bonferroni correction for multiple testing as reported in Lush et al. 2019l. **D)** t-SNE plot of *tnfb*-expressing cells from homeostatic and from neuromast cells (0-10 hours) post HC ablation. *tnfb*-expressing cells are located within the cluster of mantle cells and in some cells from the central support cell cluster. Heat bar shows log2 fold change. **E)** Heatmap of *tnfb* expression within each cell cluster and at various timepoints: one before (pre-HC) and several after HC ablation. Homeostatic mantle cells are enriched in *tnfb* mRNA, and directly after HC ablation (0h), *tnfb* transcripts increase in mantle cells. At the same timepoint, *tnfb* transcripts also increase in central support cells. **F)** Violin plot of normalised *tnfb* expression within mantle cells in homeostatic cells (pre-HC ablation) and at several timepoints post-HC ablation. *tnfb* transcripts increase early and transiently following HC ablation. Asterisks (*) indicate significant difference (p<0.001) assessed by Wilcoxon Rank test followed by a Bonferroni correction for multiple testing as reported in Beak et al. 2022.

**Supplementary Figure 4.**
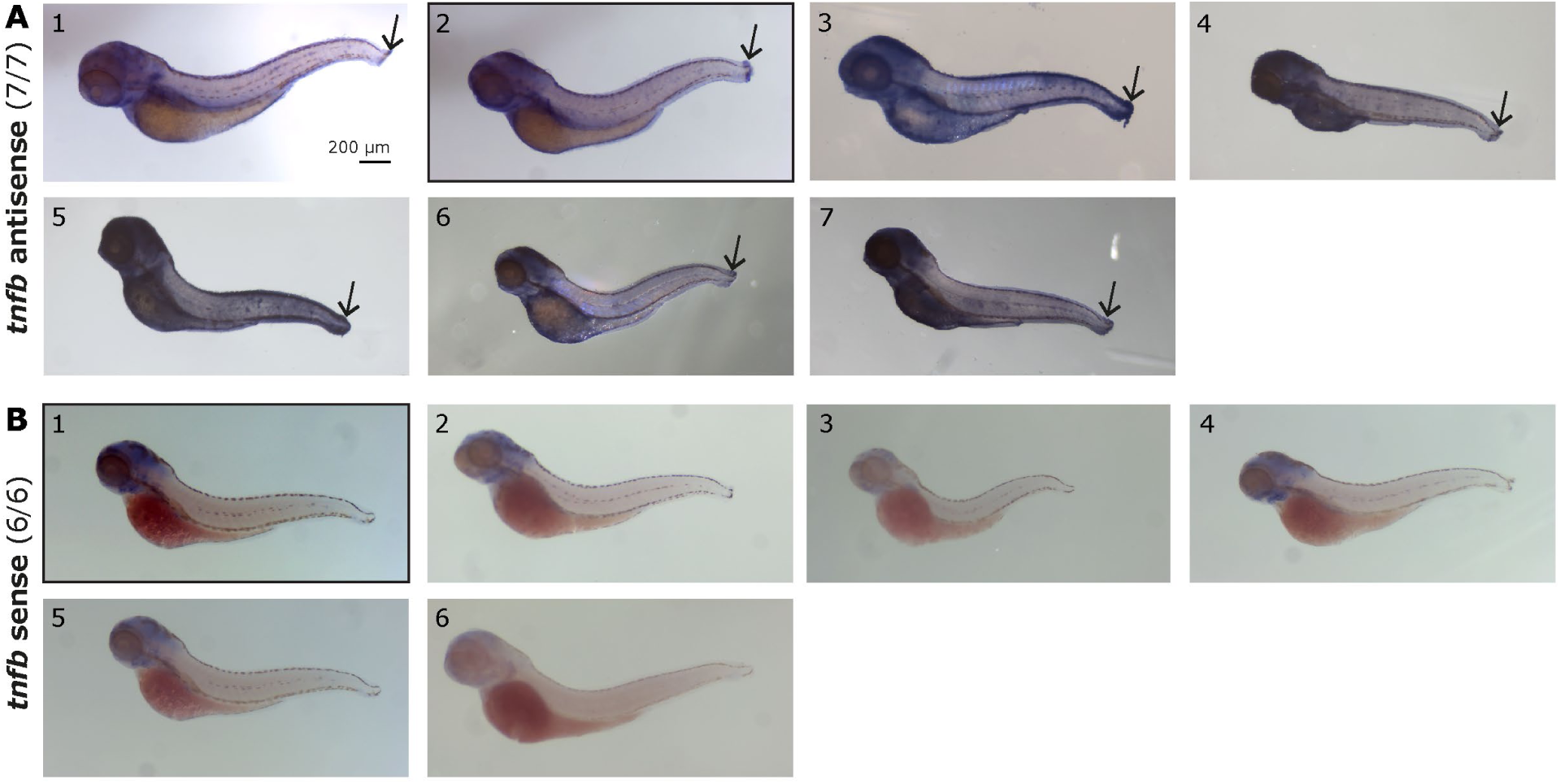
Wholemount *in situ* hybridization of *tnfb* mRNA in zebrafish after wounding. Wholemount *in situ* hybridization with *tnfb* antisense and sense probes in 3 dpf larvae, 2 hours after caudal fin fold amputation (black arrows). **A)** Panel of *tnfb* mRNA expression. Larvae were scored positive when signal was visible in the tail. **B)** *tnfb sense* probe did not show any specific signal, some coloration in the head was observed. Black outlined are the images of larvae also shown in main Figure 3B.

**Supplementary Figure 5.**
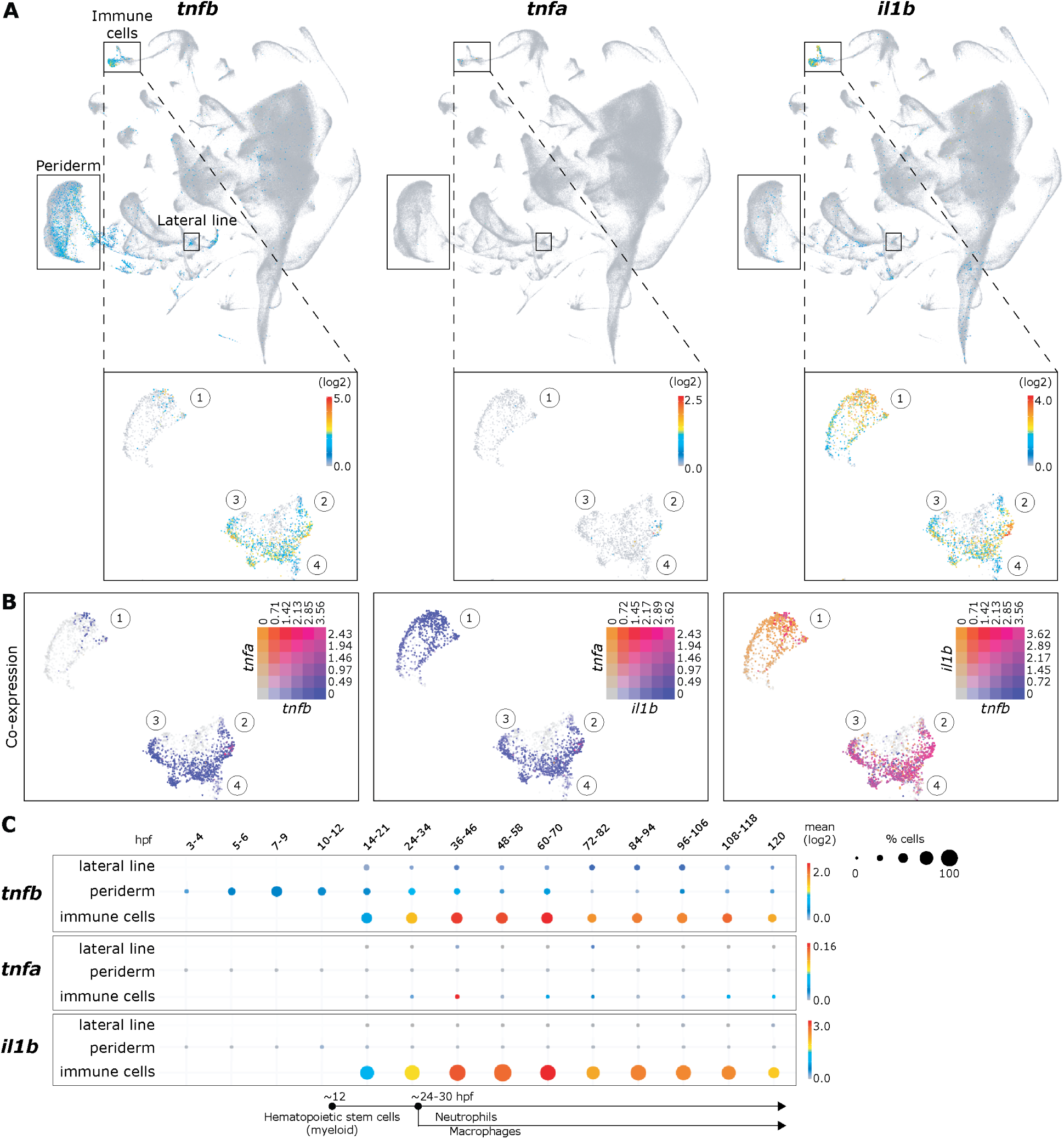
Innate immune cells of developing zebrafish are enriched in *tnfb* and *il1b,* and not *tnfa* transcripts. Single-cell RNA-sequencing performed across multiple zebrafish developmental stages (3 – 120 hpf) was retrieved from <u>Daniocell</u> database and plots were generated using the web interface (A/C) or the companion application <u>Daniocell Desktop (B)</u>. **A**) t-distributed Stochastic Neighbour Embedding (t-SNE) plots of cytokines *tnfb, tnfa,* and *il1b* in all cells, accompanied by a panel, plotted to the same scale, zoomed in on four innate immune cell clusters: 1. neutrophils, 2. macrophages, 3. microglia, 4. hematopoietic stem cell (myeloid). While *tnfb* and *il1b* are enriched at varying levels in macrophages, microglia, haematopoietic stem cells (myeloid) and in neutrophils, *tnfa* could only be detected at a low level in a few macrophages. **B**) Co-expression of *tnfb/tnfa/il1b* transcripts. Matrix indicates relative expression level per cell. Only a few macrophages are *tnfa^+^tnfb^+^* or *tnfa^+^il1b^+^*. In contrast, *tnfb^+^il1b^+^* cells are present in all four cell clusters, including in a subpopulation of neutrophils. **C**) Mean enrichment of cytokine transcripts across multiple developmental stages (3 – 120 hpf) in lateral line, periderm (non-immune tissue) and in immune cells. *tnfb* is initially detected in periderm, prior to the emergence of immune cells, and by 14-21 hpf onwards *tnfb* transcripts are enriched in lateral line and immune cells. Enrichment of *il1b* is also evident in immune cells. At the bottom, the timeline indicates the emergence of hematopoietic stem cells (myeloid), followed by differentiation and appearance of neutrophils and macrophages.

## Supplementary Videos

**Supplementary Video 1. 3D rendering of *tnfb:mCh-F*-expressing cells within a neuromast.** Representative neuromast from *Tg(tnfb:mCherry-F)* generated as 3D project in ImageJ-FIJI. The rotating projection shows that mCherry^+^ cells form a hollow dome-like shape typical of neuromast mantle cells.

**Supplementary Video 2.** *Tg*(*tnfb:eGFP-F)* and *Tg*(*tnfb:mCherry-F)* report inducible *tnfb-*expression in the same motile (immune) cells. Caudal fins of 3 dpf *Tg(tnfb:eGFP-F;tnfb:mCherry-F)* larvae were amputated or left intact (control) and images were acquired over a period of 10 hpa as a 21 µm z-stack at a 18-minutes interval with an Andor spinning disk confocal microscope at 20x magnification. Representative time-lapse of the maximum projections rendered at 5 frames per second are shown. Please note that, in control (left time-lapse) and in amputated (right time-lapse) fins, tissue displacement is observed due to growth. Neuromast are therefore used as reference point within the tissue to distinguish motile from stationary cells within the tissue, independent of tissue growth. In both time-lapse, left green arrow indicates the relative position of a eGFP^+^mCherry^-^cells to the nearest neuromast; the right green arrow indicates the relative position between two eGFP^+^mCherry^-^cells. In all cases the relative positions do not change. In the intact fin, few eGFP^+^mCherry^+^ cells are observed and move non-directionally throughout the tissue; eGFP^+^mCherry^-^cells however are stationary relative to the neuromast (green arrows). In the amputated fin, the number of eGFP^+^mCherry^+^ cells increase, and these cells are very motile, and most move directionally towards the wound; eGFP^+^mCherry^-^cells however, although they change their morphology by protruding dendrites, do not move directionally towards the wound. As the tissue regenerates and grows, eGFP^+^mCherry^-^cells appear to move towards the right, but they do not change their position relative to the neuromast (green arrows). Larvae as shown are the same as those in main Figure 3A.

**Supplementary video 3. Differential onset of *tnfb* expression in macrophages and neutrophils.** Caudal fins of 3 dpf transgenic larvae were amputated or left intact as control. Images were acquired over a period of 6 hpa as a 21 µm z-stack at a 6-minutes interval with an Andor spinning disk confocal microscope at 20x magnification. Representative time-lapse of the maximum projections, rendered at 5 frames per second, are shown. The video first shows *Tg(tnfb:mCherry-F;mpeg1:eGFP),* followed by *Tg(tnfb:mCherry-F;mpx:eGFP),* together showing that *tnfb:mCh-F* expression is detectable in macrophages earlier than in neutrophils.

**Supplementary video 4. Kinetics of expression of *tnfb, tnfa* and *il1b* after amputation.** Caudal fins of 3 dpf transgenic larvae were amputated or left intact and imaged at a 6-minute interval with an Andor spinning disk confocal microscope using 20x magnification. Representative time-lapse covering the 9 hpa period are shown, rendered at 5 frames per second. Each frame is a maximum projection of images acquired as 21 z-stacks. Tracks were added to indicate representative *tnfb:mCh-F*^+^ cells also expressing either *tnfa:eGFP-F or il1b:eGFP-F*. The video first shows *Tg(tnfa:eGFP-F);Tg(tnfb:mCherry-F),* followed by *Tg(il1b:eGFP);Tg(tnfb:mCherry-F)*.

**Supplementary video 5. Kinetics of *tnfa:eGFP-F* and *tnfb:mCh-F* expression in macrophages following *E. coli* injection.** Transgenic zebrafish (3 dpf) were injected intramuscularly with 1 nl PVP or with 2×10^3^ CFU *E. coli* crimson resuspended in PVP, and images were acquired over a period of 5 hpi as a 58 µm z-stack at a 10-minutes interval with an Andor spinning disk confocal microscope at 20x magnification. Representative time-lapse of the maximum projections rendered at 5 frames per second are shown. In the trunk of a PVP-injected *Tg(tnfb:mCherry-F;mpeg1:eGFP)* larvae, few macrophages (eGFP^+^) migrate to the site of injection and express tnfb (eGFP^+^mCherry^+^), the number of recruited macrophages expressing *tnfb* (eGFP^+^mCherry^+^) increases following to *E. coli* injection. In the trunk of a PVP-injected *Tg(tnfa:eGFP-F;mpeg1:mCherry-F)* larvae, few macrophages (mCherry^+^) are recruited to the site of injection express tnfa (eGFP^+^mCherry^+^), the number of recruited macrophages expressing tnfa (eGFP^+^mCherry^+^) increases following *E. coli* injection.

